# Easy and Accurate Reconstruction of Whole HIV Genomes from Short-Read Sequence Data

**DOI:** 10.1101/092916

**Authors:** Chris Wymant, François Blanquart, Astrid Gall, Margreet Bakker, Daniela Bezemer, Nicholas J. Croucher, Tanya Golubchik, Matthew Hall, Mariska Hillebregt, Swee Hoe Ong, Jan Albert, Norbert Bannert, Jacques Fellay, Katrien Fransen, Annabelle Gourlay, M. Kate Grabowski, Barbara Gunsenheimer-Bartmeyer, Huldrych F. Günthard, Pia Kivelä, Roger Kouyos, Oliver Laeyendecker, Kirsi Liitsola, Laurence Meyer, Kholoud Porter, Matti Ristola, Ard van Sighem, Guido Vanham, Ben Berkhout, Marion Cornelissen, Paul Kellam, Peter Reiss, Christophe Fraser, The BEEHIVE Collaboration

## Abstract

Next-generation sequencing has yet to be widely adopted for HIV. The difficulty of accurately reconstructing the consensus sequence of a quasispecies from reads (short fragments of DNA) in the presence of rapid between- and within-host evolution may have presented a barrier. In particular, mapping (aligning) reads to a reference sequence leads to biased loss of information; this bias can distort epidemiological and evolutionary conclusions. *De novo* assembly avoids this bias by effectively aligning the reads to themselves, producing a set of sequences called contigs. However contigs provide only a partial summary of the reads, misassembly may result in their having an incorrect structure, and no information is available at parts of the genome where contigs could not be assembled. To address these problems we developed the tool `shiver` to preprocess reads for quality and contamination, then map them to a reference tailored to the sample using corrected contigs supplemented with existing reference sequences. Run with two commands per sample, it can easily be used for large heterogeneous data sets. We use `shiver` to reconstruct the consensus sequence and minority variant information from paired-end short-read data produced with the Illumina platform, for 65 existing publicly available samples and 50 new samples. We show the systematic superiority of mapping to `shiver`’s constructed reference over mapping the same reads to the standard reference HXB2: an average of 29 bases per sample are called differently, of which 98.5% are supported by higher coverage. We also provide a practical guide to working with imperfect contigs.

## 1 Introduction

The genetic sequences of pathogens are a rich data source for studying their epidemiology and evolution, and provide information for vaccine and therapeutic design. In the past decade, next-generation sequencing (NGS) has transformed genomics, with decreasing costs and enormous increases in the amount of data available. Despite the success of NGS in other fields, sequencing of human immunodeficiency virus (HIV) is still largely based on the older method of Sanger sequencing. For example, on the comprehensive Los Alamos HIV database [1], of the 119,237 samples with platform information, 91.6% were generated by Sanger sequencing, 7.0% with the Roche 454 platform, 1.4% with Illumina platforms, and 0.02% with the IonTorrent platform. Restricting to the 38,635 samples dating from 2010 or later, these numbers change only to 94.6% Sanger sequencing, 2.0% 454, 3.4% Illumina and 0.02% IonTorrent.

More broadly, NGS has been hugely successful both for sequencing samples with no within-sample diversity, and at the opposite end of the spectrum, for metagenomic studies. In the first case, any apparent within-sample diversity is attributable to sequencing error; in the latter case, there is no presumption that different fragments of DNA have the same origin, and so each fragment is checked against large databases to catalogue within-sample diversity [2, 3].

HIV is an intermediate case: the long duration of chronic infection coupled with high rates of replication and mutation mean that a single infection, and hence a single sample, will contain a diverse collection of related viral particles, frequently called a quasispecies. Reconstructing different aspects of these quasispecies from *reads* (fragments of sequence; see Fig. 1) has proven technically challenging [4] and may have been a significant obstacle to the widespread adoption of NGS for HIV. Here, we present an easy to use program developed for this task. Note that a variety of NGS platforms exist, which can be broadly classified into short-read-low-error platforms and long-read-high-error platforms (see e.g. [5]); here we focus on the former.

**Figure 1:**
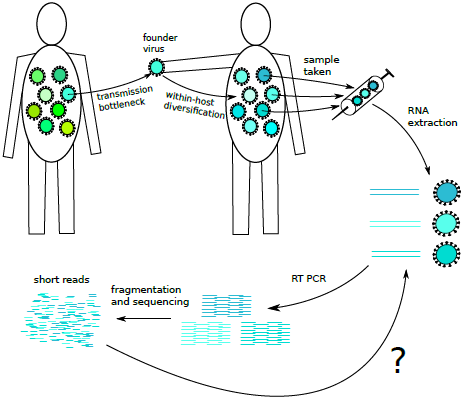
Interpreting next-generation sequencing data for HIV.

The complex problem of quasispecies reconstruction can be bypassed with single genome amplification (SGA): in SGA, by limiting dilution, samples are reduced to single-virion aliquots that are sequenced separately [6–8]. However, the costs of using SGA for large population studies would be prohibitively high. Our program was developed as part of the BEEHIVE project *(Bridging the Evolution and Epidemiology of HIV in Europe)* in which samples from over 3,000 individuals with known date of HIV infection are being sequenced to investigate the viral-molecular basis of virulence [9]. The power of genome-wide association studies, and of epidemiological analyses for example identifying transmission risk factors, is enhanced by focussing resources on the widest possible population coverage.

The quasispecies in one patient can be summarised by the consensus sequence – the ‘average’ sequence of those virions sampled, as represented in the reads. Determining the most common base at each position in the genome, and which other bases are present and at what frequencies, requires the reads to be mapped (aligned) to a reference sequence. To what should they be Mapped? Mapping to a reference too far from the quasispecies’ true consensus leads to biased loss of information [10–13]. Like any form of sequence alignment, mapping relies upon sequence similarity; the more a read differs from its reference, the less likely it is to be aligned correctly or at all. This hides differences between the sample and the reference, giving a consensus genome erroneously similar the reference chosen.

The implications of this problem for downstream sequence analysis are worrying. Using the same reference for multiple patients will tend to make their consensuses artefactually similar, overestimating proximity in a transmission network and distorting epidemiological conclusions. Using old reference sequences to construct new ones biases the new to resemble the old, which could distort our picture of evolution and hinder monitoring of emerging virulent or resistant variants. As an example, for a survey of *env* gene diversity in currently circulating viruses, it would be highly undesirable to artificially bias the reconstructed sequence towards similarity with the standard HXB2 reference virus isolated in 1983.

### 1.1 Mapping Reads: Problems and Solutions

The problem is acute in the case of RNA viruses like HIV: rapid mutation and substitution generate large within-and between-host diversity, compounding the difficulty of mapping reads accurately. Fig. 2 shows an example of biased data loss, in which an insertion in the sample is lost because it is missing in the reference to which the reads were mapped. Indeed insertions and deletions (in-dels) are very common in HIV [15,16], especially in the *env* gene [17], and reads from indels are particularly difficult to map correctly [18–21].

The loss of reads during mapping is roughly proportional to the divergence between the true consensus and the reference used, with for example 10% divergence giving a 25% loss of reads [11]. In the *env* gene, divergence of greater than 10% can arise in the course of a single infection [22–24]. Crucially, the loss of data is also biased, with the bias occurring at different scales. Data is more likely to be lost in (i) those samples in a dataset that differ more greatly from the reference used for their mapping; (ii) those parts of the genome, in a single sample, where the sample and reference are most different; and (iii) a subset of genotypes, in a single diverse sample, that are more different from the reference than the other genotypes.

**Figure 2:**
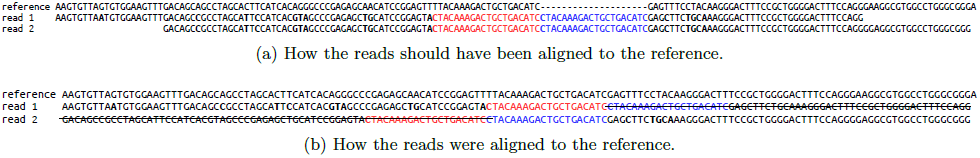
An example of biased loss of information encountered in our data when mapping to an existing reference. The reads contain a 20 base pair (bp) duplication – the sequence shown first in red then again in blue — which the reference B.JP.05.DR6538.AB287363 does not have. Correct alignment, shown in the upper panel, would have inserted a 20bp gap into the reference to accommodate the duplication. What the mapper actually did (lower panel) was to either align the first occurrence of the 20bp sequence to its match in the reference and discard everything after it (read 1), or align the second occurrence and discard everything before it (read 2). ‘read 1’ and ‘read 2’ each represent thousands of similar reads; their consensus is therefore well supported but misses the duplication. This bias occurred despite the reference being the same subtype as the sample (B), and having been singled out by the program kraken [14] as the closest of 160 references to this set of reads.

This problem means the simplest mapping strategy – using as a reference some existing, standard genome that is expected to be similar to the sample has much room for improvement. For example one could map once to a standard reference, call the consensus, then use this as the reference for one or more rounds of remapping [13, 25–28]. Remapping is expected to be more accurate, because the consensus initially called is expected to be closer to the true consensus than the standard reference is. For this to be the case all along the genome however, reads must map correctly all along the genome in the first step. If the sample has an indel not present in the reference, inaccurate mapping at the site of the indel may cause it to be missed when the consensus is called, as in Fig. 2. Remapping is then doomed to repeat the same error.

To correct for this, between initial mapping to the standard reference and calling the first consensus, multiple sequence alignment can be performed with the reads [11, 29]. In this case reads do not need to map *correctly* all along the genome, since realignment should correct misalignment around indels, but they do still need to *map* all along the genome. If biased data loss leads to a failure of reads to map at a given point, the missing reads will not shape the initial consensus and remapping to that consensus will not recover them. For the variable loop regions of HIV’s *env* gene in particular, reads from one virus can easily fail to map to another; many examples of this can be seen in Appendices F and G, manifest as genomic windows in which reads do map to a reference tailored to the sample, but not to the standard HIV reference HXB2, resulting in missing sequence in the latter case. (As specific examples see the V1-V2 loop region in Figures 39, 49 50, 56 and 57.)

These problems motivate *de novo* assembly (hereafter just assembly). Roughly, this consists of aligning overlapping reads to each other, tolerating some pre-set level of disagreement between them to allow for some within-sample diversity or sequencing error, iteratively extending using reads overhanging the edges, finally resulting in aset of sequences called contigs (see e.g. [30]). Remapping to contigs [12, 13, 26, 31, 32] settles ambiguity at positions spanned by multiple contigs which disagree, corrects positions where assembly did not call the most common base, provides minority variant information, and allows greater use to be made of base quality information than is typically done during assembly.

However, contigs may differ from the true consensus by more than just a few single nucleotide polymorphisms (SNPs), which are easily corrected by mapping. Misassembly may occur, giving contigs supported by a high depth of reads but whose structure is very different from the known genome. This can arise *in silico* [13], i.e. by misassembly of correct reads; or as a result of chimeric reads produced during sequencing, due to recombination during library preparation [13,33,34], concatemerisation/ligation [35], or stem loops of RNA secondary structure [32].

Furthermore, the set of contigs resulting from assembly may not fully cover the genome. Gaps between contigs can be due to a total absence of reads there, following sequencing failure or only a partial genome present in the sample. They can also be due to the reads being too few (though non-zero), or too diverse, for successful assembly; in this case, mapping can recover consensus sequence not present in assembly output.

Finally, as the set of reads will generally contain contamination, so will the set of contigs. These should be identified and discarded.

To address these problems we developed the tool `shiver` – *Sequences from HIV Easily Reconstructed* – to preprocess and map reads from each sample to a custom reference, tailored to be as close as possible to the expected consensus, constructed by correcting contigs and filling in gaps between them with the closest identified existing reference sequences. We wrote it to be easy to use, suitable for simple scripted application to large heterogeneous data sets, in this population genomics study and elsewhere.

## 2 Results

For HIV samples sequenced with the Illumina platform yielding paired-end short read data, we produced consensus sequences, together with summary minority-variant information (base frequencies at each position) and detailed minority-variant information (all reads aligned to their correct position in the genome). Our tool `shiver` also produces a single alignment containing all of the consensuses separately generated for each sample. All resulting sequence data will be deposited in public repositories on publication of this preprint. The input data constituted 68 samples previously sequenced with Miseq, and 50 samples newly sequenced with Hiseq (see Methods). Only 65 of the Miseq samples had contigs that returned a BLASTN hit to a sequence in our existing HIV reference set; these and all 50 Hiseq samples were fully processed, giving whole or partial genomes.

Appendices F and G contain figures showing the genes of HIV in their reading frames, a set of sequences, and the coverage (number of reads mapped at each position) along the genome, for each sample. We reproduce the figure for the first Miseq sample here – Fig. 3 – as an example for discussion. The sequences shown are, from top to bottom: the standard reference sequence HXB2, the reference created and used for mapping by `shiver`, the consensus of reads mapped to `shiver` reference, the consensus of reads mapped to HXB2 (the exact same reads, i.e. following `shiver`’s removal of adapters, primers and low quality bases; then mapped with identical parameters), and the contigs generated by *de novo* assembly. One the pa-rameters of `shiver` is the minimum coverage required to call the base at each position; here we chose 10, since the assembler we used – `IVA` – requires at least 10 reads to extend a contig. Vertical black lines inside sequences in the alignment denote SNPs, defined here relative to the most common base among these sequences. Horizontal black lines indicate a lack of bases, i.e. a deletion relative to another sequence in the alignment or, for the two consensuses, simply missing sequence due to coverage being less than 10. In Fig. 3 we see an amplification failure confined to the region around the vif and vpr genes: no contig sequence, a coverage less than 10, and so no consensus sequence. The information contained in the few reads that did map to this region is retained in the minority variant files produced by `shiver`; consensus sequence could be called here, if one chose to lower the minimum coverage threshold parameter below 10.

**Figure 3:**
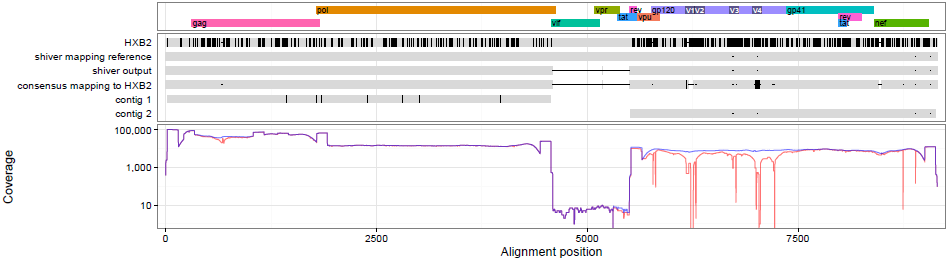
Top panel: genes in their reading frames. Middle panel: sequences for the Miseq sample ERR732065. From top to bottom these are the standard reference sequence HXB2, the reference created and used for mapping by `shiver`, the consensus of reads mapped to `shiver` reference, the consensus of the same reads mapped to HXB2, and the contigs. Bottom panel: the coverage (number of mapped reads) for the `shiver` reference in blue, and for HXB2 in red.

Where HXB2 and the sample differ by many close SNPs or an indel, differences between the shiver and HXB2-derived consensuses arise. The coverage plot beneath the sequences shows that at such points, the coverage mapping to HXB2 almost always drops below the coverage mapping to the `shiver` reference; given that the same reads are being mapped to the same part of the genome, this strongly suggests that the `shiver` consensus is more accurate. For example in Fig. 3, in the *gag* gene, HXB2 has a deletion relative to the sample. Mapping to HXB2 then results in a consensus erroneously containing this deletion, with a local drop in coverage. (Though coverage drops, it is still more than 10,000, showing that a large absolute number of reads is no guarantee of accuracy.) Mapping to the `shiver` reference on the other hand does not introduce the deletion, and the coverage remains smooth. Similar errors mapping to HXB2 can be seen in Fig. 3 in *vpu*, the four variable loops V1-V4, and *nef*.

Comparisons of these sequences are quantified for each sample in Appendix E, and in summary in Tables 1 and 2. For example Table 1 shows that mapping one sample’s reads to `shiver`’s reference instead of HXB2 gives a mean increase in consensus length of 136bp, and a mean number of 29 bases called differently. This latter figure is broken down by coverage with respect to the two references the appendix, with the result that at 98.5% of these disagreeing positions, the `shiver` reference has higher coverage. Interpreting higher coverage as more accurate mapping, mapping to the `shiver` reference instead of HXB2 corrects 28:7 false SNPs and introduces 0:4 false SNPs per sample.

**Table 1:** Comparing the consensus from mapping to the reference constructed by `shiver` with the consensus from mapping to HXB2. Minima, medians, means and maxima are over the combined set of 65 Miseq and 50 Hiseq samples processed. Medians and means are rounded to the nearest integer.

|  |  |  |
| --- | --- | --- |
| Number of bases differing between the two consensuses: | min | 0 |
|  | median | 22 |
|  | mean | 29 |
|  | max | 177 |
| Increase in consensus length from mapping to the **shiver** reference instead of HXB2: | min | 0 |
|  | median | 118 |
|  | mean | 136 |
|  | max | 581 |

Table 2 shows that the `shiver` consensus is on average 144 bases longer than the set of contigs. The mean number of bases in the `shiver` consensus that differ from all contigs at point is 14. This last figure is biased favourably towards the contigs, however, as the comparison was made after `shiver` performed contig correction. This is because a comparison of two sequences requires them to be aligned, and aligning the *spliced* or partially reverse-complemented contigs that `shiver` corrects (see Methods) would give a nonsensical alignment.

**Table 2:** Comparing the consensus from mapping to the reference constructed by `shiver` with the corrected contigs. Minima, medians, means and maxima are over the combined set of 65 Miseq and 50 Hiseq samples processed. Medians and means are rounded to the nearest integer.

|  |  |  |
| --- | --- | --- |
| Number of bases differing between the consensus and all contigs at that point: | min | 0 |
|  | median | 7 |
|  | mean | 14 |
|  | max | 106 |
| Length of sequence present in contigs but missing from the consensus: | min | 0 |
|  | median | 0 |
|  | mean | 0 |
|  | max | 1 |
| Length of sequence present in the consensus but missing from the contigs: | min | 6 |
|  | median | 38 |
|  | mean | 144 |
|  | max | 2445 |

These differences are small compared to the length of the HIV genome: approximately 9000 bases. However the aim of sequencing a known pathogen is not to produce a roughly correct picture of the known genome, but to obtain each sample’s sequence as accurately as possible, so that the number of differences between similar samples can be meaningfully interpreted.

Among the reads mapped by `shiver`, interesting within-host diversity is maintained, capable of revealing structure in the quasispecies. Fig. 4 shows an example for our Hiseq sample 17796_3_29. The reads are from the boundary between p2 and p7 in the *gag* gene; roughly a third of them have a 21bp insertion relative to the others. This insertion is not seen in any other sequence in the Los Alamos National Laboratory alignment HIV1_ALL_2015_gag._DNA’ of 7903 *gag* sequences [1]. Though not a duplication at the nucleotide level, it duplicates the *GATAMMQ* amino acid motif. Mutations at the p2/p7 boundary [36] and insertions at other *gag* cleavage sites [37] have been implicated in restoring replicative capacity in viruses treated with protease inhibitors.

**Figure 4:**
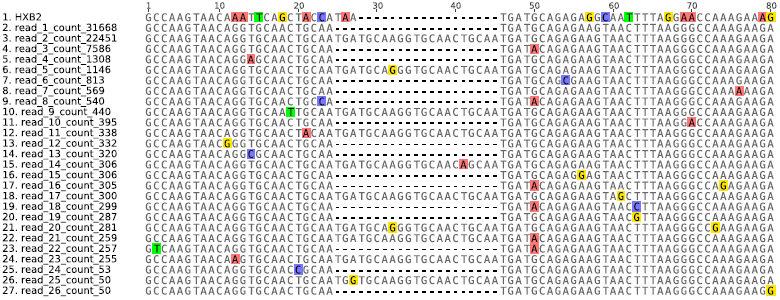
Within-host indel polymorphism in our Hiseq sample 17796_3_29: a 21bp insertion in roughly a third of the reads duplicates the *GATAMMQ* amino acid motif at the boundary between p2 and p7 in *gag*. The value following ‘_count_’ in the sequence name is the number of times that exact sequence was found in the reads here following mapping with `shiver`; only sequences found at least 50 times are shown. HXB2 is included for comparison. Coloured squares highlight bases differing from the consensus; bases without a coloured square agree with the consensus base at that position (ignoring gaps).

## 3 Methods

### 3.1 Data Summary

The 68 Miseq samples we considered were those sequenced and released with the `IVA` publication [38], namely accession numbers ERR732065–ERR732132. The short reads were downloaded from the European Nucleotide Archive.

The 50 Hiseq samples we considered were newly generated for the BEEHIVE project, from confirmed seroconverters from Europe. RNA was extracted manually from blood samples following the procedure of [39]. This was amplified using universal primers that define four overlapping amplicons spanning the whole genome, following the procedure of [40], then sequenced.

For both sets of samples short reads were assembled into contigs using `IVA`, which has been shown to outperform other viral assemblers for HIV [38].

See Appendix A for more details.

### 3.2 Method Summary

The steps in our method `shiver` are shown in Fig. 5; see Appendix B for more details.

**Figure 5:**
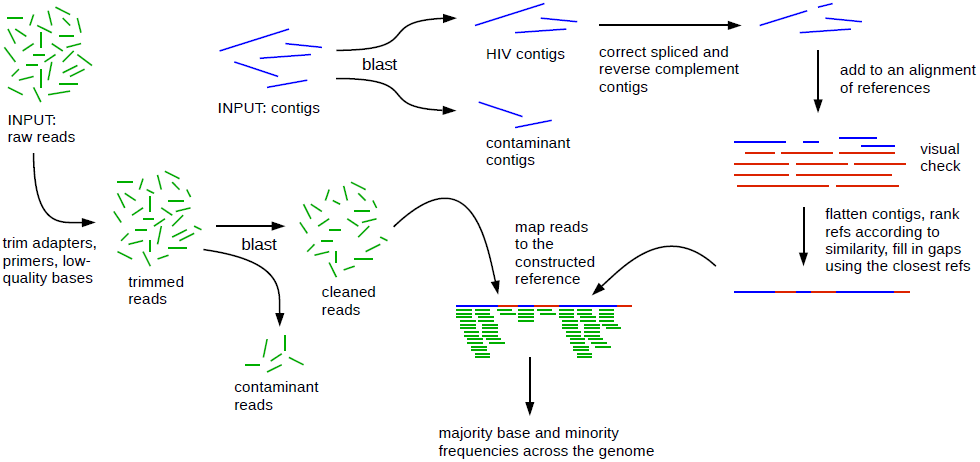
A summary of the steps in our method `shiver`.

In summary: paired-end short reads and contigs assembled from those reads are required as input. Comparison to a set of existing reference genomes separates the contigs into those that are HIV and those that are contaminant. Spliced contigs – those concatenating two separated regions of the genome into a single sequence – are cut, then any contigs in the opposite orientation to the existing references are reverse-complemented. The motivation for this cutting of contigs is the assumption that HIV does not exhibit major structural variation, e.g. variation in gene presence/absence or gene order, which is supported by sequence compendiums to date [1]. The contigs are added to the alignment of existing references. Here `shiver` stops to allow a visual check of the correctness of this alignment. Once it is checked, `shiver` continues (all remaining steps in the program are performed by with the second of two commands needed for full processing). The alignment of contigs to existing references is used to create a reference for mapping which is tailored to the sample. This is done by using contig sequence where available, and those existing references that match the contigs most closely to fill in any gaps between contigs. Before mapping, reads are trimmed for low-quality bases, adapter and primer sequences; contaminant read pairs are diagnosed as those matching contaminant contigs more closely than the tailored reference, and are removed. The remaining reads are mapped to the tailored reference, each position in the genome is considered (resolving indel polymorphism) to find the frequencies of different bases, and the most common base is called to give the consensus. Optionally, the cleaned reads can be remapped to the consensus (with missing coverage in the consensus filled in with the corresponding part of the tailored reference), for a second iteration of calling the base frequencies and the consensus. This was done for the data processed here (which explains why the `shiver` reference does not match the contigs exactly in Fig. 3 and the figures of Appendices F and G).

`shiver` also produces a ‘global alignment’ of all consensuses it generates by coordinate translation, without need for an alignment algorithm, including correct placement of partial genomes split into segments separated by missing coverage.

### 3.3 Running shiver Fully Automatically

Alternatively `shiver` can be run from beginning to end without the break in the middle described above, for applications where visually checking the contigs is impractical. This is only possible for samples not requiring contig correction, and does not produce the global alignment of all samples’ consensuses together. The different alignment strategy used in this case, and our recommendation that that the contigs be checked instead, are discussed further in Appendix B.

#### 3.4 Using shiver

`shiver` and its documentation are available at github.com/ChrisHIV/shiver. It was designed to be run in Linux-like environments, including Mac OS. Once dependent packages are installed, `shiver` itself requires no installation: it is a set of executable scripts. The Genomic Virtual Laboratory [41], provided for example on the UK Medical Research Centre’s Cloud Infrastructure for Microbial Bioinformatics (MRC CLIMB) [42], contains all dependencies^1^, allowing `shiver` to be run immediately.

Before processing with `shiver`, short reads must be assembled into contigs^2^. `shiver` is run from the command line using three commands: a one-off initialisation command, then two commands per sample to be processed. This produces, for each sample,

- the mapped reads in BAM format [45]
- a plain text file with the counts of the different bases at each position;
- the consensus;
- a coordinate-translated version of the consensus for a global alignment;
- an alignment of the consensus and the contigs, for comparison;
- the insert-size distribution; and
- a separate BAM file of only the contaminant reads mapped to the HIV reference (illustrating the importance of removing these reads prior to mapping).

The global alignment is constructed simply by combining the coordinate-translated consensus files from all samples into one file, e.g. running from the command line `cat file1 file2 […] > MyGlobalAlignment. fasta`

For our data, `shiver` typically look less than an hour to process each Miseq sample, and up to ten hours for each Hiseq sample (the latter containing roughly ten times as many reads).

All bioinformatic parameters can be changed in a configuration file, allowing customisation of how reads are trimmed, how they are mapped, and how the consensus is called as a function of coverage and diversity. `shiver` also includes simple command-line tools for partial reprocessing (modifying sample output without rerunning the whole pipeline), and for analysis: see Appendix D.

#### 4 Discussion

We developed the tool `shiver` to preprocess and map reads from each sample to a custom reference, constructed using *de novo* assembled contigs supplemented by existing reference genomes. Tailoring the reference to be as close as possible to the expected consensus before mapping maximises the accuracy of the mapping, and therefore of the resulting consensus. `shiver`’s identification, ranking, and use of the closest existing references to fill in gaps between contigs boosts data recovery for samples with amplification failure or assembly failure. Such partial-genome samples, which are inevitable in large diverse data sets, are processed with exactly the same two commands; this simplifies scripted application of shiver to all samples in a data set. `shiver` also produces a global alignment containing all of the consensuses separately generated for each sample, which is usually required for comparative analysis of the sequences such as for phylogenetics or genome wide association studies (GWAS).

Mapping to `shiver’s` constructed reference instead of mapping the same reads to the standard reference HXB2 gives a consensus sequence which is on average 136 bases longer, with 29 of the original bases called differently, of which 98.5% are supported by higher coverage. This shows the importance of tailoring the reference to the sample before mapping. `shiver`’s consensus, obtained by mapping reads to a reference constructed from the contigs, has on average 14 bases called differently from the contigs even after correcting structural problems in the contigs. This highlights the need for mapping in addition to assembly.

A limitation of the method is that after reads have been successfully mapped (which makes requirements on base quality and good alignment to the reference), we consider each read to carry equal weight in determining the consensus and the frequency of variant bases. The frequency of a variant in the reads and its frequency in the sampled virions may differ due to PCR bias – amplification of some virions more than others. Reconciling these frequencies would require modelling the number of virions in the sample, their diversity, the process generating PCR bias, and sequencing error, which is beyond the scope of this work. Alternatively this problem can be addressed with the sequencing protocol: primer IDs [46] can associate every read to its template, allowing identification of all PCR duplicates (as well as permitting separate reconstruction of all haplotypes). As with single genome amplification however, higher costs for each sample currently limit applicability to large population studies.

Another limitation is that no mapping of diverse reads can guarantee perfect accuracy at every position in every sample, as perfect sequence alignment is an unsolved problem. In particular where samples contain indel polymorphisms, or where localised misassembly results in an indel not present in the reads, mapping may misalign reads in a way that is not cured by remapping to their own consensus, since the misalignment gives an error in the consensus. As with all automatic sequence alignment, there is scope for improvement by manual inspection.

A design choice is that `shiver` does not take into account translation to amino acids, and in particular does not bias towards maintaining reading frames. Deliberately including this bias would be clearly justified for many organisms, but the case is arguable for HIV due to overlapping reading frames, frame-shifting polymorphisms, and possibly antisense expression [47, 48]. Other tools exist to extract in-frame gene sequences from `shiver` consensuses, such as `Gene Cutter` [49].

Individuals who are dually infected – hosting two distinct quasispecies, whether by two distinct founder viruses establishing productive infections, or by superinfection – are known to be special cases clinically, and perhaps for evolution, because of the opportunity for recombination. It is important to note that they are also special cases for bioinformatic processing. If one of the two quasispecies is more highly represented in the reads at every position in the genome, the consensus sequence for the patient will be simply the consensus of the more abundant quasispecies. However if one quasispecies has more reads at part of the genome and the other has more reads elsewhere in the genome, the consensus will be a recombinant of both quasispecies; a recombinant which may never have existed *vivo*, and which may invalidate phylogenies in which it is included. Clearly, care must be taken in identifying such patients as their dually infected status may not be known.

Our focus here has been reconstruction of the consensus sequence that summarises a quasispecies. The process of doing this from diverse reads – from different virions in the quasispecies – retains rich information on within-host diversity. Our separate tool `phyloscanner` (Wymant, Hall *et al*., in preparation) allows easy extraction, processing, alignment and parallel phylogenetic analysis of the short reads from many genomic windows of many BAM files, for example those produced by `shiver`. Examination of within-host and between-host diversity together, at every position along the genome, allows identification of dual infections, transmission, recombination and contamination. These more detailed pictures of quasispecies and the relationships between them, in addition to their summaries as consensus sequences, further motivate the valuable role next-generation sequencing has to play in our understanding of HIV.

## Acknowledgements

Thanks to Martin Hunt, Dan Frampton and Tiziano Gallo Cassarino for helpful discussions, and to Simon Burbidge and Matt Harvey for help with Imperial College London High Performance Cluster computing. This work was funded by ERC Advanced Grant PBDR-339251. This work used the computing resources of the UK MEDical BlOinformatics partnership – aggregation, integration, visualisation and analysis of large, complex data (UK MED-BIO) which is supported by the Medical Research Council [grant number MR/L01632X/1].

## Footnotes

1 Except `smalt`, which is loaded on MRC CLIMB with the single command `brew install smalt`, and otherwise available at [43].

2 This important step, though difficult technically, it is not difficult for the user: our chosen assembler `IVA` can be run on a virtual machine provided by the Sanger pathogens group [44], and assembles contigs from reads with a single command from the command line. The user can use any assembler; others are available in the Genomic Virtual Laboratory, including `SPAdes, Velvet` and `MIRA`, though currently none designed for viral data.

## Appendix A Our Data in More Detail

*The existing Miseq data.* The short reads for these samples were publicly released with [38]. The samples have different origins; six are from a longitudinally sampled transmission pair studied in [50]. ERR732065-ERR732072 were sequenced with 150bp reads, ERR732073-ERR732132 with 250bp reads. Note that only 42 of these 68 samples were assembled in [38]: the rest failed quality control checks designed to pre-select robust whole-genome samples. We reassembled all 68 samples with `IVA` for processing with `shiver`, as by design the method can be run in exactly the same way for those samples devoid of genuine sequence, those with partial genomes and those with whole genomes.

*The new Hiseq data.* 5*μ*l of amplicon 1 (the shortest and most successfully amplified amplicon) was pooled with 10*μ*l each of amplicons 2-4. Multiple samples were pooled during library preparation, using one of 192 multiplex adaptors for each sample. The library was sequenced in ‘rapid run mode’ on both lanes of a HiSeq2500 instrument with read lengths of 2 × 250bp, resulting in two lanes of short reads per sample. Automatic processing at the Wellcome Trust Sanger Institute used `IVA` to generate contigs for each lane, i.e. two sets of contigs per sample. We combined the two sets to allow comparison of the assembly output resulting from two technical replicates of short reads. For the large majority of cases the contigs were nearly identical, but stochastic differences in the read populations between lanes mean the resulting contigs occasionally differ. Insert sizes were typically less than 500bp, i.e. mates in a pair overlap.

The 50 Hiseq samples were chosen from a larger data set currently being collected and sequenced for the BEEHIVE project’s primary aim of investigating the viral-molecular basis of virulence. Selection criteria for inclusion in the project include a known date of infection, either by negative and positive tests separated by less than a year, or by clinical signs of acute infection at diagnosis; and a sample obtained for sequencing between 6 and 24 months after diagnosis, before beginning antiretroviral treatment and before progression to AIDS. The 50 samples processed here were chosen as follows. (i) One sample chosen with a large difference in the fraction of the genome assembled between the two Hiseq lanes, as an example of the variability of assembly output. (ii) Nine samples chosen with misassembled contigs for one or both Hiseq lanes, to illustrate the necessity of `shiver`’s contig correction. (iii) From each of the Dutch, French, German and Swiss cohorts, ten samples with contigs spanning the whole genome: five with a `COMET` [51] subtype result of unambiguously pure subtype B, five with a result of unambiguously non-B or ambiguous.

## Appendix B Our Method in More Detail

An alignment of existing reference sequences is required as input. Construction of a custom reference for mapping involves identifying the existing references that are closest to the sample under consideration. The greater the number and diversity of existing references given as input, the denser the coverage of sequence space is and the closer the closest reference is expected to be, with corresponding benefits for the accuracy of the results. However these existing references should be aligned to each other accurately, in order for the addition of each sample’s contigs to the alignment to be meaningful; this means that producing such an input by simply automatically aligning many diverse sequences is not advised. This alignment will be used as input for every sample processed by `shiver`, and so careful manual curation is time well spent. We used the 2012 ‘Compendium’ alignment from the Los Alamos National Laboratory HIV database [52], which is already manually curated.

Custom reference construction begins with contig pre-processing as follows. Matches between the contigs and any existing reference from the alignment are searched for using `BLASTN` [53] with default settings. Contaminant sequence is inevitable in high-throughput NGS; any contig that has no BLASTN hit to any of the HIV references is taken to be contamination, and is put aside for later use, leaving contigs that are putatively HIV. Assembly may produce contigs that are erroneously spliced – concatenating two separated regions of the genome – due to errors *in silico* or during sequencing as mentioned. We detect such misassemblies from multiple `BLASTN` hits for a single contig (discarding any hit wholly contained inside another hit), and correct them by cutting. After cutting, any contig sequence in the opposite orientation is reverse-complemented. (`IVA` outputs contigs such that the longest open reading frame is on the forward strand.) Contigs are then aligned to the existing reference alignment using `MAFFT` [54], trying both `‑‑add` and `‑‑addfragments` modes and using the one with the smallest maximum gap fraction (the maximum calculated over all contigs in each alignment).

The alignment of contigs to the set of existing references should be visually inspected at this point. For HIV sequences, reference [16] states that “Algorithmic alignment does not necessarily retrieve the best alignment. It is important to always verify whether the sequence data are aligned unambiguously and, if necessary, manually correct the alignment.” [13] echoes this for any evolving pathogen: “the ‘best’ alignment chosen by an alignment program is not necessarily the ‘true’ alignment… Alignment quality should also be inspected manually in a visualisation program”. The commonness of indels in HIV [15, 16] compound the problem of misalignment. As well as revealing alignment error, inspecting the aligned contigs allows detection of problems with the contigs themselves, further discussed in Appendix C. In our experience such problems are confined to the minority of samples; nevertheless the importance of finding them for downstream analysis, and the ease of correcting them, motivates checking every sample. We used `Geneious` [55] for sequence visualisation and editing.

Using the alignment of contigs to existing references, the set of contigs is flattened into a single sequence as follows. At positions covered by one contig, its base (or gap character, for a deletion) is used. At positions covered by multiple contigs, if all contigs have a gap we use a gap, else only contigs with a base here are considered: if all contigs agree on the base we use that base; else the contigs disagree on the base, and we use that of the longest contig. We used this ‘base of the longest contig’ heuristic expecting that, where sufficiently distinct haplotypes exist to result in multiple contigs covering the same place, haplotypes supported by a higher depth of reads would tend to be assembled into longer contigs. The heuristic’s suitability was confirmed posthoc by seeing that at positions where more than one base was seen in different contigs, 78% of the time the `shiver` consensus agreed with the base of the longest contig. The heuristic’s importance is minimal, since at only 0.5% of positions where one contig had a base did a second contig have a different base. These figures refer to the combined Miseq and Hiseq data presented here; more diverse samples or different assemblers could both result in more disagreement between contigs.

The sequence resulting from this flattening of the contigs is compared to each existing reference in the alignment in turn, counting identical bases including gaps within contigs (known deletions) but not gaps between contigs (missing information), allowing a ranking by similarity. As existing references have variable lengths (the long terminal repeat regions that flank the clinical genome are often not sequenced; genes may terminate prematurely), the closest reference is extended outwards using any overhanging sequence from the second closest reference, then the third longest sequence etc. terminating when both edges of the alignment are reached. This sequence – the elongated closest reference – is used to fill in any gaps between (but not inside of) the flattened contigs. This completes production of the reference tailored for this sample.

Before mapping to this reference, the reads are trimmed and cleaned as follows. Adapters, primers and low quality bases are trimmed using `Trimmomatic` [56] and `Fastaq` [57]. We then consider contaminant reads from non-HIV sources. Most of these would presumably be discarded by mapping to an HIV reference, due to lack of similarity. However there is ample opportunity for traces of human DNA to end up in a sample, and sequence of endogenous retroviruses in human DNA may resemble HIV. As a guard against this, and other contamination resembling HIV, we use `BLASTN` to find all read pairs that are a better match to one of the contigs previously found to be contamination, than to the tailored reference. These pairs are discarded.

The cleaned reads are mapped to the tailored reference with `SMALT` [43], giving a file in BAM format. Using `SAMtools` [45] the BAM file is read into pileup format, which is parsed to give base frequencies at each position in the genome. Note that within-host diversity does not consist exclusively of point mutations: indels can be present in some reads and not others(Fig. 4 is an example), which must be dealt with in the pileup. Where some reads have a deletion relative to the reference and others do not, the deletion/gap character can simply be considered as a fifth base whose frequency can be counted like the others. Where some reads have an insertion relative to the reference and others do not, or more generally where insertions of two or more sizes are present, we find the most common insertion size and, inside that insertion, consider only those reads with an insertion of that size (thus avoiding any ambiguity in the alignment of the inserted sequences to each other). Finally, the base frequency file is parsed to call the consensus base at each position, with thresholds on coverage and the diversity required for ambiguity codes.

Since we know how the consensus aligns to the reference used for mapping, and we know how that reference (constructed from the contigs) aligns to the input alignment of existing references, we can construct a global alignment of the consensuses from all samples merely by coordinate translation, negating the need for further alignment and manual curation. Two things must be excised from the consensus for this global alignment reconstruction: insertions present in the majority of reads but not in their tailored reference (which are rare, since the reference is constructed from the contigs which are constructed from the reads), and insertions present in the contigs but none of the existing references (which are rare provided the set of existing references is large and diverse). In both cases this is sequence whose alignment to the common anchor of the existing references is not known, and so coordinate translation cannot align it.

As mentioned, `shiver` can be run from beginning to end without the break in the middle, with the single command `shiver_full_auto.sh`, for uses where visually checking the contigs is impractical. This begins with separation of contigs into HIV (those with `BLASTN` hits) and contamination as previously. Subsequent steps are as follows.

1. The need for contig correction is checked, but correction is not performed: if it is needed, processing stops. Trust in the accuracy of an automated alignment of contigs cut into pieces based on evidence of structural problems would be trust misplaced.
2. Each HIV contig is now certain to have a single `BLASTN` hit (discarding any smaller hits wholly contained inside others). That hit is checked to span some minimum fraction of the contig length (by default 90%) as a guard against contigs containing containing some foreign sequence; otherwise processing stops.
3. Multiple sequence alignment is performed with these contigs and just one of the existing reference sequences, for each of the existing reference sequences separately.
4. For each such alignment, generated both with regular `mafft` and with `mafft --addfragments`, we calculate the fractional agreement between the flattened contigs and the reference, i.e. the fraction of positions spanned by the reference and at least one contig where the reference and the longest contig agree. Misalignment is penalised in this score because gaps inside contigs are taken as genuine deletions.
5. For the alignment with the highest score, the maximum gap fraction amongst the contigs in the alignment (i.e. the fraction of positions inside the contig that are gaps) is checked to be below a user-specified threshold (the default is 5%, based on analysis of thousands of such alignments that we visually checked) as a further guard against misalignment.
6. The contigs are flattened using this single existing reference to fill in any gaps between them, generating the mapping reference tailored for this sample.

Aligning contigs to the references one at a time (step 3) is simpler for the alignment algorithm than aligning to all of them at once, and means that even if misalignment occurs for what is truly the closest reference to the contigs, the alignment to the second closest can be used instead. Trimming of low-quality bases, trimming of adapter and primer sequences, removal of contaminant reads and mapping to the tailored reference all occur as described previously. For samples that cannot be processed fully automatically this way – when contig correction is required, or a contig is spanned by too small a `BLASTN` hit, or too many gaps are present after alignment – the main mode of `shiver` is available (requiring inspection of the contigs).

As argued earlier, we advocate visually inspecting the aligned contigs, i.e. running the two-command implementation of `shiver` (with the check in between commands). This also has the advantage of working for all samples, whereas shiver_full_auto.sh will not proceed if problems with the contigs or their alignment are detected. shiver_full_auto.sh also does not produce a global alignment of all consensuses to each other, because the coordinate translation procedure allowing its construction is derived from each sample’s alignment of contigs to all of the references at once. That alignment is produced for two-command implementation of `shiver`, but step 3 above aligns contigs to references one at a time.

## Appendix C Working with Imperfect Contigs

The generation of perfect contigs from real, diverse, imperfect short read data, for every sample in an arbitrarily large data set, is an unsolved problem. One could discard any contig that is automatically detected to be suspicious in some respect; however this discards valuable information. More pragmatically, one can look at the contigs. Specifically, inspecting an alignment of one sample’s contigs with a diverse set of existing reference sequences allows one to judge whether the pattern of SNPs and indels in the contigs is consistent with the diversity amongst the references. What problems with the contigs can be detected and corrected when inspecting such an alignment?

- False-positive ‘HIV’ contigs: designated as HIV by virtue of having a `BLASTN` hit to the existing references, but very poorly aligned to the existing references, throughout the contig's length. Poor alignment means implausibly many SNPs and/or implausibly many indels and/or implausibly large indels, relative to the existing references. Such contigs are probably assembled from non-HIV contaminant reads, with just enough similarity to have a `BLASTN` hit; they should simply be discarded.
- Localised misassembly at a contig end: the ends of contigs are by definition points at which the assembler has been unable continue extending the sequence, due either to lack of reads, or to diversity too high for a sensible representative sequence to be chosen. The latter possibility also means erroneous bases are sometimes called in short stretches of sequence at the end of a contig, which align poorly. Trimming such sequence from the ends of contigs means the corresponding sequence from the closest existing reference will be used instead, giving a better reference for mapping. (Some assembly algorithms trim the ends from contigs for precisely this reason; however the trimming may not be sufficient.)
- Structurally misassembled contigs: those that are spliced, concatenating two or more separated regions of the genome. `shiver` corrects these by cutting between the regions, allowing for their independent alignment. When cut correctly, no action from the user is needed. However, such contigs are detected by having two (or more) `BLASTN` hits, neither contained wholly inside the other; this can also arise from an unusual but genuine indel. A judgement call is then needed – whether the indel is a misassembly or real, whether the contig should be cut or not – which can be informed by considering indels in other existing reference sequences at this point. With long enough reads and sufficiently accurate mapping, reads will map here correctly whether or not the reference constructed from the contigs contains the indel, making the question moot; however with mapping inaccuracies of the kind shown in Fig. 2 possible, it’s best to get the reference’s structure as correct as possible before mapping.
- Contigs wholly or partially reverse-complemented, relative to the set of reference sequences to which one wants to align. If the assembler does not orientate the contigs, on average half of them will be in the reverse orientation; `IVA` orientates contigs such that the longest open reading frame is on the forward strand, however for very short contigs this may fail. Contigs wholly in the reverse orientation simply need to be wholly reverse-complemented. In the process of assembling a spliced contig, an assembler may concatenate different regions in different orientations; each region may or may not require reverse-complementation after cutting into separate regions. `shiver` does this; no contig correction in this respect should be required by the user.

## Appendix D Sample Reprocessing and Analysis

Individual steps from `shiver` can be run with stand-alone command line tools, for ease of reapplication elsewhere. For example `CorrectContigs.py` is run with a file of contigs and a file of their `BLASTN` hits, and corrects the contigs by cutting and reverse-complementing where needed. Also included in `shiver` are command-line tools for easy analysis and modification of sample output without rerunning the whole pipeline:

- Two parameters specified in the configuration file are a minimum coverage required to call a base (below this coverage, the character ‘`?`’ is used) and a larger minimum coverage required to use upper case instead of lower, as an easy signal of increased confidence. (Note that decreasing these parameters will, in general, allow bases to be called at more positions, giving a longer consensus. However there is a trade-off: where there are fewer reads, the effect of contaminant reads on the consensus may be greater.) To regenerate a consensus with new values of these parameters, `CallConsensus.py` can be run on a sample’s base frequencies file. To regenerate a coordinate-translated version of this consensus for the global alignment (of all consensuses produced by `shiver`), `TranslateSeqForGlobalAln.py` can be run on the consensus.
- Another parameter in the configuration file is the minimum read *identity* – the fraction of bases in the read which are mapped and agree with the reference – required for a read to be considered mapped, and so retained in the BAM file. If you wish to increase this after completion of `shiver`, reads with an identity below your new higher threshold can be discarded by running `RemoveDivergentReads.py` on a BAM file. Running shiver_reprocess_bam.sh on the resulting BAM file (or indeed any BAM file) implements just the last steps in `shiver`, namely generating pileup, calculating the base frequencies, and calling the consensus.
- `FindNumMappedBases.py` calculates the total number of mapped bases in a BAM file (where read length is constant this equals. the number of mapped reads multiplied by read length, minus the total length of sequence clipped from reads), optionally binned by read identity. In the absence of mapped contaminant reads, and all else being equal, mapping to a reference which is closer to the true consensus should map more bases and mapped reads should have higher identities.
- `FindClippingHotSpots.py` counts, at each position in the genome, the number and percentage of reads that are clipped from that position to their left or right end. Having many such reads is a warning sign of the kind of biased loss of information shown in Fig. 2b.
- `LinkIdentityToCoverage.py` finds, for each different coverage encountered when considering all positions in a BAM file, the mean read identity at such positions. The mean read identity tends to be lower at positions of low coverage due to a background of contaminant reads, which differ from the reference by virtue of being contamination, but which are nevertheless similar enough to be mapped. Quantifying the decline in identity at low coverage helps inform what coverage threshold is appropriate for a given data set.
- `AlignMoreSeqsToPairWithMissingCoverage.py` allows more sequences to be added to a pairwise alignment in which one sequence contains missing coverage (such as a consensus and its reference), correctly maintaining the distinction between gaps (indicating a deletion) and missing coverage.
- `AlignBaseFreqFiles.py` aligns not two sequences, but two of the csv-format base frequency files output by `shiver`. Optionally a similarity metric is calculated at each position in the alignment, between 0 (no agreement on which bases/gaps are present) and 1 (perfect agreement on which bases/gaps are present and on their proportions). This allows comparison not just of consensus sequences between two samples but also of minority variants.
- `ConvertAlnToColourCodes.py` converts each base in a sequence alignment into a colour code indicating agreement with the consensus and indels; `AlignmentPlotting.R` takes such colour codes and visualises the alignment. These two scripts were used to produce the plots of Appendices F and G.
- Finally some simple tools for convenience: `FindSeqsInFasta.py` extracts named sequences from a fasta file, with options including gap stripping, returning only windows of the sequences, and inverting the search; `PrintSeqLengths.py` prints sequence lengths with or without gaps; `SplitFasta.py` splits a fasta file into one file per sequence therein.

## Appendix E Sequence Statistics

The final column of Tables 3 and 4 – the length of sequence present in the `shiver` consensus but absent from the contigs – has the value 38 for many samples. This happens because the combined length of the first and last amplification primers is 38. Reads ending in a perfect match to one of primers (or their reverse complements) have that end trimmed, both during assembly of the reads into contigs by `IVA`, and as part of `shiver` before mapping the reads. Occasional sequencing error results in a base in the primer being miscalled, meaning a small fraction of reads containing the primer do not have it trimmed. Because the error is random, a variety of slightly different variations on the true primer are seen. This collection of variant sequences is too diverse to be assembled: when extending a contig by adding a given kmer to its end, `IVA` requires that the kmer be at least four times as abundant as the next most abundant kmer. `IVA` therefore stops assembling the contig at the primer’s edge (i.e. not including the primer). When mapping reads at this position however, as each base miscall is random, a meaningful consensus of the variants can be called, namely that of the true primer sequence. This phenomenom suggests that differences in sequences lengths between the contigs and the consensus which are ≲ 38 do not represent meaningful increases in genome coverage obtained by `shiver` mapping compared to the contigs.

**Table 3:** Comparison of different sequences resulting from processing the 65 Miseq samples.

| Sample | shiver<br>consensus<br>length | Comparing [the consensus mapping to the **shiver**<br>reference] to [the consensus mapping to HXB2]. |  |  | Comparing [the consensus mapping to the<br>**shiver** reference] to [the corrected contigs]. |  |  |
| --- | --- | --- | --- | --- | --- | --- | --- |
|  |  | (**shiver** con-<br>sensus length) -<br>(HXB2 consen-<br>sus length) | Number of dis-<br>agreeing bases<br>where the **shiver**<br>consensus has a<br>higher coverage | Number of dis-<br>agreeing bases<br>where the HXB2<br>consensus has a<br>higher coverage | Number of po-<br>sitions disagree-<br>ing between the<br>consensus and<br>all contigs | length of<br>sequence only<br>in<br>contigs | length of se-<br>quence only in<br>**shiver** consensus<br>(see note on next<br>page) |
| ERR732065 | 8236 | 142 | 3 | 0 | 7 | 0 | 87 |
| ERR732066 | 7371 | 118 | 0 | 0 | 0 | 0 | 35 |
| ERR732067 | 5784 | 94 | 14 | 2 | 5 | 0 | 868 |
| ERR732068 | 4578 | 2 | 0 | 0 | 3 | 0 | 39 |
| ERR732069 | 5733 | 55 | 3 | 1 | 20 | 0 | 60 |
| ERR732070 | 8115 | 223 | 54 | 2 | 28 | 0 | 71 |
| ERR732071 | 8259 | 265 | 6 | 0 | 13 | 0 | 68 |
| ERR732072 | 8126 | 58 | 27 | 5 | 40 | 0 | 79 |
| ERR732073 | 9120 | 255 | 66 | 2 | 67 | 0 | 38 |
| ERR732074 | 9085 | 241 | 63 | 0 | 44 | 0 | 38 |
| ERR732076 | 9091 | 289 | 23 | 0 | 4 | 0 | 38 |
| ERR732077 | 9091 | 250 | 16 | 0 | 0 | 0 | 38 |
| ERR732078 | 9091 | 249 | 32 | 0 | 0 | 0 | 38 |
| ERR732079 | 9091 | 303 | 25 | 0 | 7 | 0 | 38 |
| ERR732080 | 9067 | 240 | 36 | 0 | 9 | 0 | 38 |
| ERR732081 | 9073 | 267 | 38 | 0 | 8 | 0 | 38 |
| ERR732082 | 9055 | 472 | 35 | 0 | 13 | 0 | 17 |
| ERR732083 | 9064 | 49 | 24 | 0 | 15 | 0 | 38 |
| ERR732085 | 9034 | 62 | 50 | 8 | 44 | 0 | 1954 |
| ERR732086 | 9040 | 47 | 44 | 0 | 13 | 0 | 38 |
| ERR732087 | 9040 | 61 | 42 | 0 | 4 | 0 | 38 |
| ERR732088 | 9065 | 176 | 81 | 0 | 25 | 0 | 38 |
| ERR732089 | 9043 | 47 | 57 | 0 | 13 | 0 | 38 |
| ERR732090 | 9040 | 53 | 59 | 5 | 22 | 0 | 38 |
| ERR732091 | 9086 | 254 | 55 | 0 | 29 | 0 | 38 |
| ERR732092 | 9068 | 209 | 47 | 0 | 28 | 0 | 38 |
| ERR732093 | 7329 | 100 | 3 | 0 | 2 | 0 | 2445 |
| ERR732094 | 9063 | 131 | 40 | 0 | 52 | 0 | 38 |
| ERR732095 | 9058 | 147 | 76 | 0 | 106 | 0 | 38 |
| ERR732096 | 9063 | 124 | 96 | 0 | 36 | 0 | 155 |
| ERR732097 | 9127 | 268 | 58 | 0 | 61 | 0 | 38 |
| ERR732098 | 9046 | 152 | 29 | 0 | 2 | 0 | 26 |
| ERR732099 | 9058 | 142 | 9 | 0 | 1 | 0 | 38 |
| ERR732100 | 9068 | 349 | 26 | 0 | 14 | 0 | 38 |
| ERR732101 | 9074 | 140 | 43 | 1 | 32 | 0 | 38 |
| ERR732102 | 9088 | 201 | 57 | 0 | 47 | 0 | 38 |
| ERR732103 | 9034 | 63 | 48 | 0 | 14 | 0 | 188 |
| ERR732104 | 9067 | 228 | 40 | 0 | 10 | 0 | 38 |
| ERR732105 | 9083 | 268 | 47 | 0 | 13 | 0 | 38 |
| ERR732106 | 8999 | 46 | 18 | 0 | 54 | 0 | 38 |
| ERR732107 | 9005 | 131 | 8 | 0 | 7 | 0 | 32 |
| ERR732108 | 9048 | 92 | 10 | 0 | 5 | 0 | 32 |
| ERR732109 | 4575 | 0 | 1 | 1 | 2 | 0 | 32 |
| ERR732110 | 9075 | 310 | 8 | 0 | 18 | 0 | 38 |
| ERR732111 | 9102 | 401 | 8 | 0 | 37 | 0 | 38 |
| ERR732112 | 8165 | 120 | 7 | 0 | 3 | 0 | 55 |
| ERR732113 | 7370 | 136 | 13 | 0 | 6 | 0 | 35 |
| ERR732114 | 9046 | 128 | 14 | 0 | 37 | 0 | 38 |
| ERR732115 | 7974 | 171 | 15 | 0 | 9 | 0 | 643 |
| ERR732116 | 9037 | 43 | 12 | 0 | 25 | 0 | 38 |
| ERR732117 | 5021 | 369 | 9 | 0 | 9 | 0 | 68 |
| ERR732118 | 9088 | 581 | 14 | 0 | 13 | 0 | 18 |
| ERR732119 | 7372 | 247 | 6 | 0 | 10 | 0 | 35 |
| ERR732120 | 7368 | 97 | 12 | 0 | 6 | 0 | 27 |
| ERR732121 | 7368 | 77 | 11 | 0 | 0 | 0 | 35 |
| ERR732122 | 7344 | 155 | 2 | 0 | 5 | 0 | 170 |
| ERR732123 | 7803 | 199 | 11 | 0 | 8 | 0 | 48 |
| ERR732124 | 1928 | 0 | 0 | 0 | 0 | 0 | 40 |
| ERR732126 | 7363 | 69 | 0 | 0 | 3 | 0 | 34 |
| ERR732127 | 1927 | 0 | 0 | 0 | 30 | 0 | 40 |
| ERR732128 | 7362 | 61 | 1 | 0 | 7 | 0 | 1255 |
| ERR732129 | 9061 | 78 | 9 | 0 | 1 | 0 | 38 |
| ERR732130 | 9080 | 106 | 11 | 0 | 0 | 0 | 38 |
| ERR732131 | 9060 | 88 | 22 | 0 | 21 | 0 | 38 |
| ERR732132 | 9058 | 57 | 25 | 0 | 23 | 0 | 38 |

**Table 4:** Comparison of different sequences resulting from processing the 50 Hiseq samples.

| Sample | shiver<br>consensus<br>length | Comparing [the consensus mapping to the **shiver**<br>reference] to [the consensus mapping to HXB2]. |  |  | Comparing [the consensus mapping to the<br>**shiver** reference] to [the corrected contigs]. |  |  |
| --- | --- | --- | --- | --- | --- | --- | --- |
|  |  | (shiver con-<br>sensus length) -<br>(HXB2 consen-<br>sus length) | Number of dis-<br>agreeing bases<br>where the **shiver**<br>consensus has a<br>higher coverage | Number of dis-<br>agreeing bases<br>where the HXB2<br>consensus has a<br>higher coverage | Number of po-<br>sitions disagree-<br>ing between the<br>consensus and<br>all contigs | length of<br>sequence<br>only in<br>contigs | length of se-<br>quence only in<br>**shiver** consensus<br>(see note below) |
| 17621.3.80 | 9058 | 144 | 93 | 1 | 22 | 0 | 38 |
| 17653.3.25 | 9045 | 60 | 26 | 0 | 6 | 0 | 38 |
| 17653.3.36 | 8970 | 6 | 14 | 0 | 3 | 0 | 173 |
| 17653.3.56 | 9031 | 101 | 56 | 0 | 6 | 0 | 38 |
| 17653.3.62 | 9085 | 141 | 70 | 0 | 28 | 0 | 38 |
| 17653.3.64 | 9095 | 146 | 83 | 0 | 9 | 0 | 38 |
| 17653.3.72 | 9044 | 62 | 9 | 0 | 3 | 0 | 38 |
| 17653.3.74 | 9069 | 68 | 15 | 0 | 2 | 0 | 38 |
| 17654.3.46 | 9064 | 42 | 16 | 0 | 11 | 0 | 21 |
| 17654.3.71 | 9079 | 125 | 53 | 1 | 6 | 0 | 38 |
| 17654.3.72 | 9109 | 89 | 25 | 0 | 2 | 0 | 38 |
| 17654.3.78 | 9084 | 78 | 11 | 0 | 8 | 0 | 38 |
| 17795.3.40 | 9091 | 69 | 26 | 8 | 8 | 0 | 38 |
| 17796.3.1 | 9114 | 122 | 32 | 0 | 3 | 0 | 35 |
| 17796.3.29 | 9012 | 60 | 33 | 4 | 17 | 0 | 2224 |
| 17796.3.30 | 8783 | 12 | 13 | 0 | 2 | 0 | 1445 |
| 17796.3.35 | 9079 | 96 | 21 | 1 | 4 | 0 | 986 |
| 18209.3.31 | 9086 | 105 | 110 | 0 | 23 | 0 | 179 |
| 18209.3.36 | 9022 | 67 | 19 | 0 | 5 | 0 | 22 |
| 18209.3.38 | 9041 | 36 | 18 | 0 | 5 | 0 | 21 |
| 19561.3.127 | 9082 | 163 | 19 | 1 | 5 | 0 | 38 |
| 19562.3.109 | 9079 | 187 | 38 | 0 | 3 | 0 | 38 |
| 19562.3.2 | 9052 | 30 | 18 | 0 | 10 | 0 | 38 |
| 19562.3.30 | 9009 | 147 | 24 | 1 | 11 | 0 | 38 |
| 19562.3.31 | 9056 | 139 | 13 | 0 | 3 | 0 | 32 |
| 19562.3.46 | 8922 | 49 | 11 | 1 | 3 | 0 | 15 |
| 19562.3.50 | 9095 | 162 | 7 | 0 | 5 | 0 | 38 |
| 19562.3.51 | 9078 | 73 | 11 | 0 | 7 | 0 | 38 |
| 19562.3.6 | 9064 | 96 | 14 | 0 | 3 | 0 | 18 |
| 19893.3.71 | 9056 | 386 | 23 | 0 | 3 | 0 | 38 |
| 19960.3.11 | 9070 | 56 | 8 | 0 | 9 | 0 | 38 |
| 19960.3.116 | 9044 | 134 | 6 | 0 | 6 | 0 | 28 |
| 19960.3.119 | 9089 | 119 | 7 | 0 | 12 | 0 | 29 |
| 19960.3.12 | 9018 | 70 | 22 | 0 | 0 | 0 | 38 |
| 19960.3.146 | 9065 | 103 | 32 | 0 | 60 | 0 | 38 |
| 19960.3.15 | 9016 | 118 | 41 | 0 | 2 | 0 | 38 |
| 19960.3.16 | 9075 | 69 | 6 | 0 | 4 | 0 | 26 |
| 19960.3.17 | 9067 | 82 | 22 | 0 | 4 | 0 | 30 |
| 19960.3.18 | 9058 | 49 | 22 | 0 | 22 | 0 | 38 |
| 19960.3.22 | 9059 | 283 | 175 | 2 | 4 | 1 | 6 |
| 19960.3.28 | 8988 | 125 | 44 | 0 | 6 | 0 | 38 |
| 19960.3.40 | 9024 | 61 | 3 | 0 | 3 | 0 | 38 |
| 19960.3.44 | 9031 | 124 | 17 | 0 | 4 | 0 | 29 |
| 19960.3.49 | 9043 | 77 | 17 | 0 | 13 | 0 | 38 |
| 19960.3.6 | 9000 | 96 | 49 | 1 | 3 | 0 | 38 |
| 19960.3.70 | 9039 | 142 | 106 | 0 | 8 | 0 | 38 |
| 19960.3.9 | 9076 | 79 | 25 | 0 | 6 | 0 | 38 |
| 20004.3.146 | 8989 | 43 | 36 | 0 | 10 | 0 | 38 |
| 20004.3.155 | 9055 | 139 | 16 | 1 | 2 | 0 | 35 |
| 20004.3.56 | 9032 | 81 | 9 | 0 | 5 | 0 | 27 |

## Appendix F Sequences and Coverage by Sample: Miseq Data

For each sample we show an alignment of the reference created and used for mapping by `shiver`, the consensus of reads mapped to this reference, the standard reference sequence HXB2, the consensus of reads mapped to HXB2 (the exact same reads, i.e. following `shiver` preprocessing, mapped with all the same parameters), and the contigs. The contigs shown are those after any correction by `shiver`, since when misassembly gives partial reverse complements, alignment gives a mess; and after manual correction where needed. The coverage (number of reads) resulting from mapping to each reference is shown with blue (`shiver` reference) and red (HXB2) lines below the alignment. For both consensus sequences a minimum coverage of 10 was required to call the base at each position, since `IVA` requires a minimum of 10 overhanging reads to extend a contig. Vertical black lines inside sequences in the alignment denote SNPs (relative to the most common base amongst the sequences here). Horizontal black lines indicate a lack of bases, i.e. a deletion relative to another sequence in the alignment or, for the two consensuses, simply missing sequence due to coverage being less than 10. Above each alignment are the genes of HIV in their respective reading frames.

**Figure 6:**
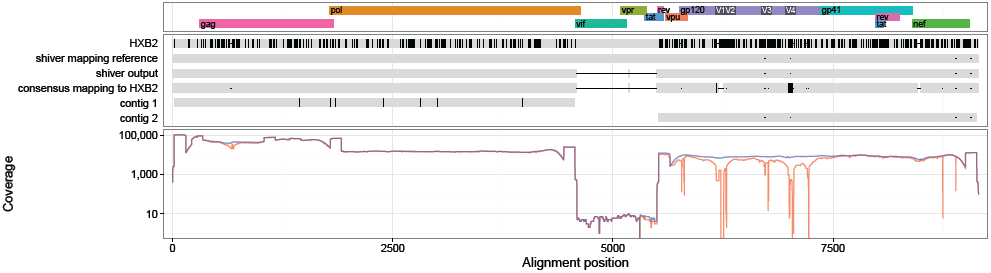
ERR732065 sequences and coverage (mapping to the `shiver` reference in blue, to HXB2 in red).

**Figure 7:**
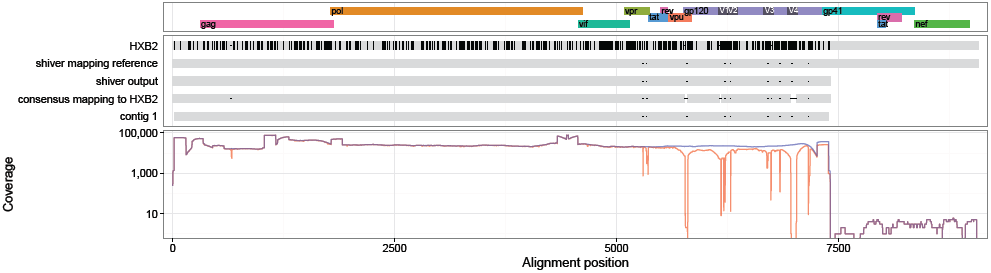
ERR732066 sequences and coverage (mapping to the `shiver` reference in blue, to HXB2 in red).

**Figure 8:**
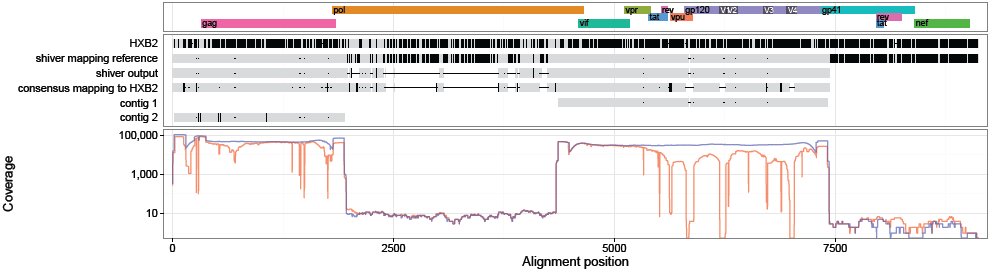
ERR732067 sequences and coverage (mapping to the `shiver` reference in blue, to HXB2 in red).

**Figure 9:**
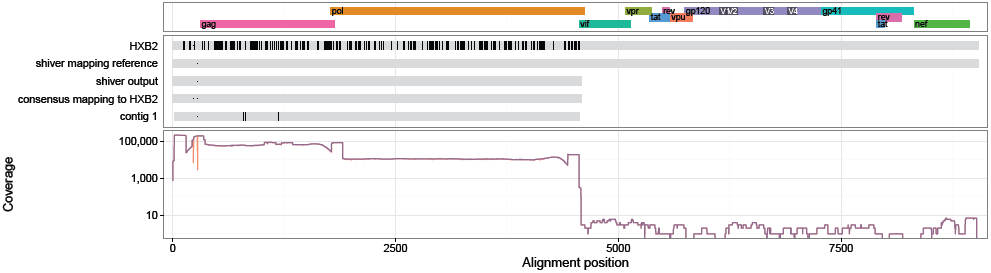
ERR732068 sequences and coverage (mapping to the `shiver` reference in blue, to HXB2 in red).

**Figure 10:**
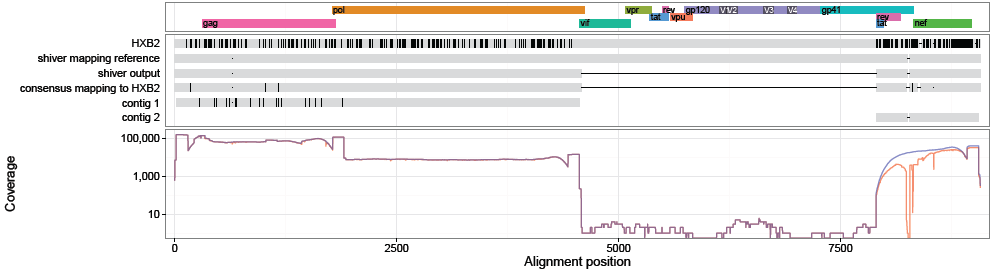
ERR732069 sequences and coverage (mapping to the `shiver` reference in blue, to HXB2 in red).

**Figure 11:**
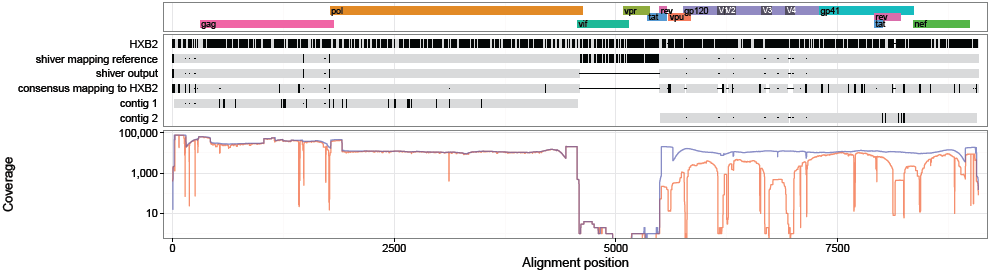
ERR732070 sequences and coverage (mapping to the `shiver` reference in blue, to HXB2 in red).

**Figure 12:**
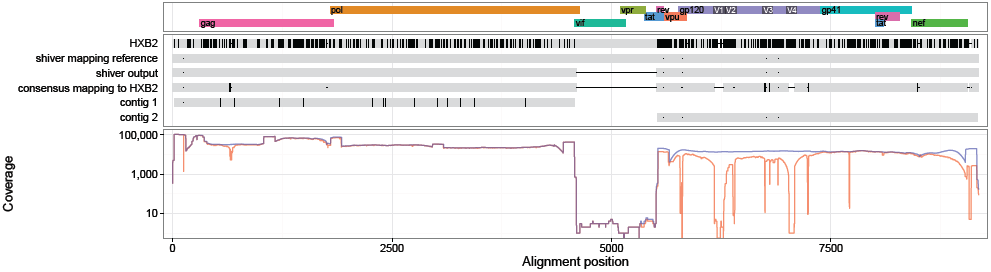
ERR732071 sequences and coverage (mapping to the `shiver` reference in blue, to HXB2 in red).

**Figure 13:**
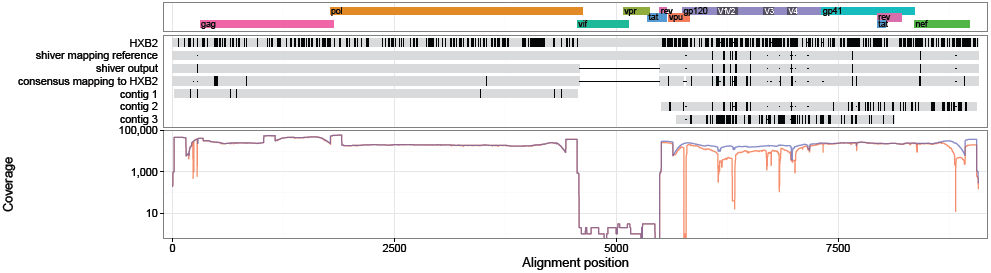
ERR732072 sequences and coverage (mapping to the `shiver` reference in blue, to HXB2 in red).

**Figure 14:**
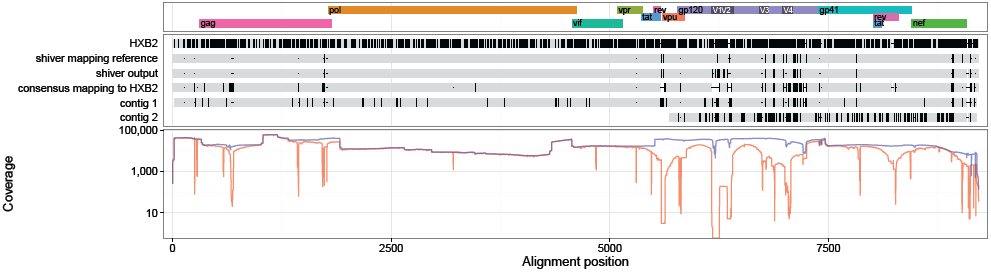
ERR732073 sequences and coverage (mapping to the `shiver` reference in blue, to HXB2 in red).

**Figure 15:**
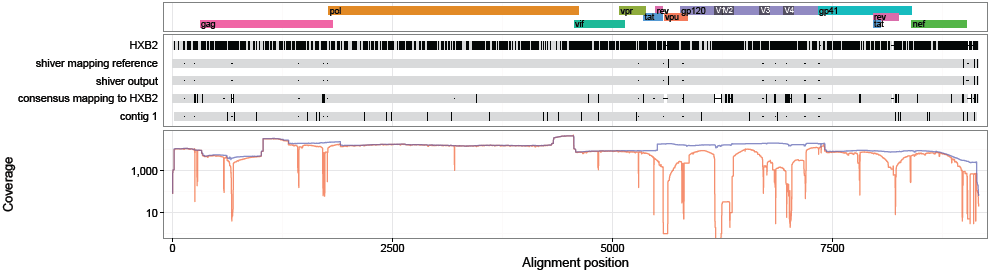
ERR732074 sequences and coverage (mapping to the `shiver` reference in blue, to HXB2 in red).

**Figure 16:**
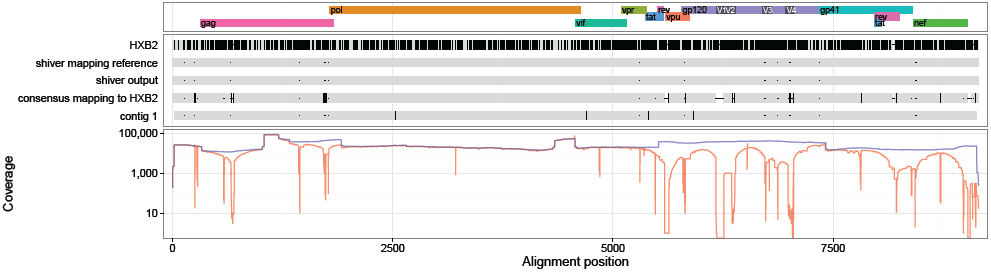
ERR732076 sequences and coverage (mapping to the `shiver` reference in blue, to HXB2 in red).

**Figure 17:**
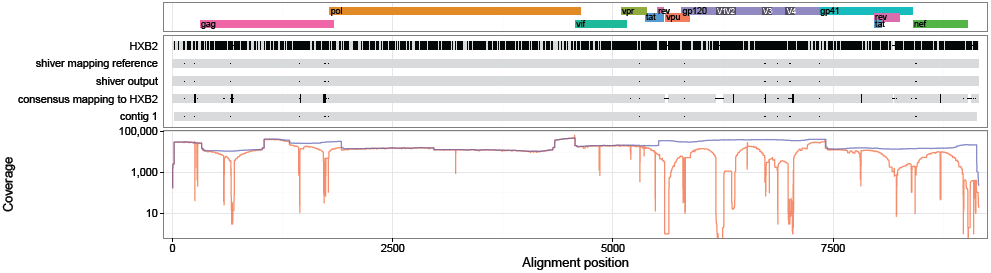
ERR732077 sequences and coverage (mapping to the `shiver` reference in blue, to HXB2 in red).

**Figure 18:**
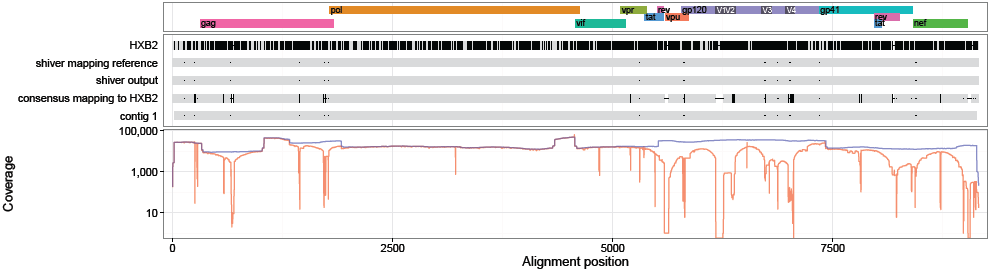
ERR732078 sequences and coverage (mapping to the `shiver` reference in blue, to HXB2 in red).

**Figure 19:**
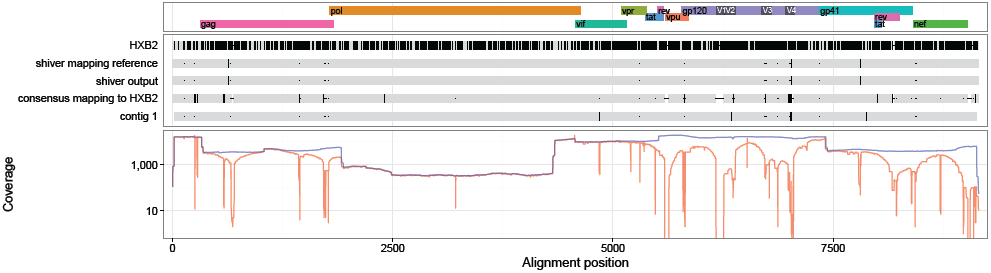
ERR732079 sequences and coverage (mapping to the `shiver` reference in blue, to HXB2 in red).

**Figure 20:**
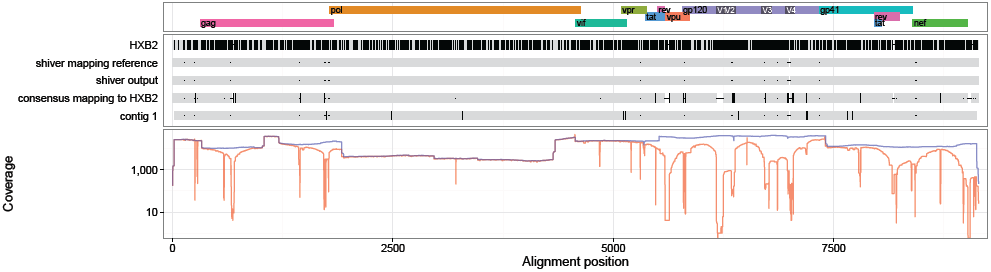
ERR732080 sequences and coverage (mapping to the `shiver` reference in blue, to HXB2 in red).

**Figure 21:**
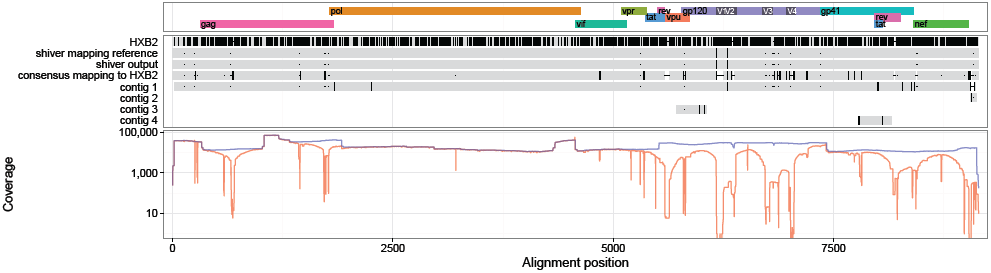
ERR732081 sequences and coverage (mapping to the `shiver` reference in blue, to HXB2 in red).

**Figure 22:**
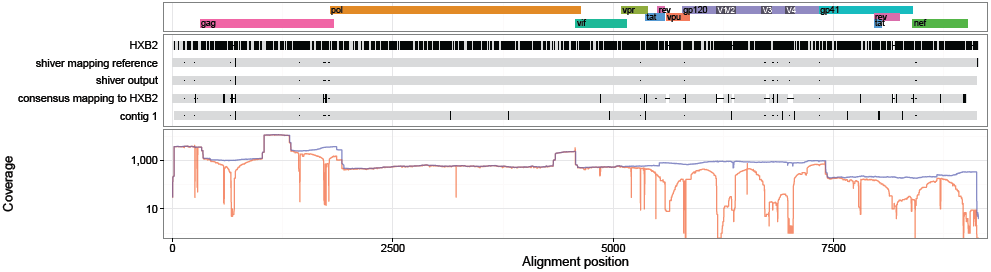
ERR732082 sequences and coverage (mapping to the `shiver` reference in blue, to HXB2 in red).

**Figure 23:**
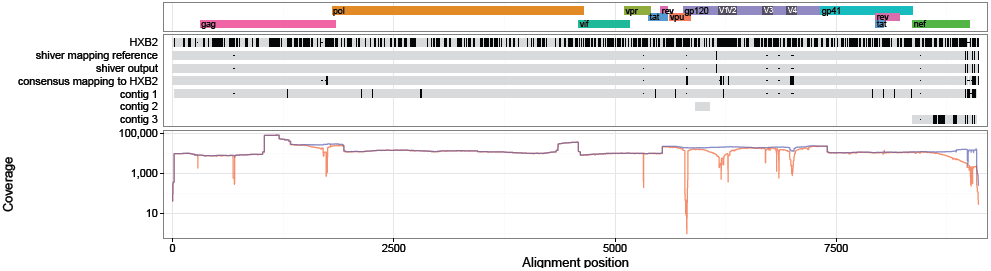
ERR732083 sequences and coverage (mapping to the `shiver` reference in blue, to HXB2 in red).

**Figure 24:**
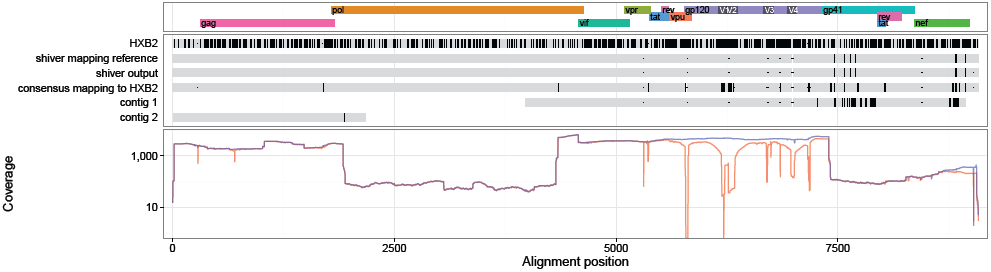
ERR732085 sequences and coverage (mapping to the `shiver` reference in blue, to HXB2 in red).

**Figure 25:**
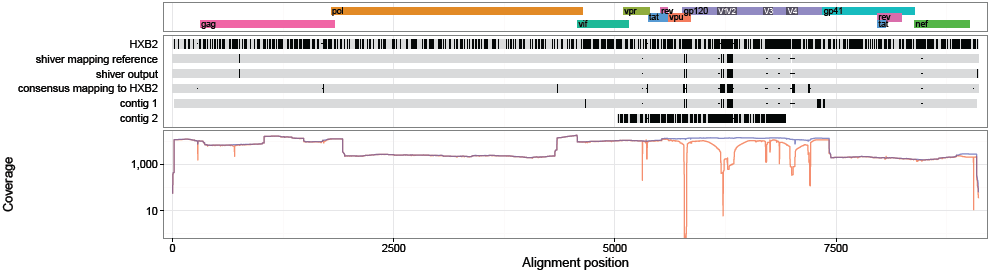
ERR732086 sequences and coverage (mapping to the `shiver` reference in blue, to HXB2 in red).

**Figure 26:**
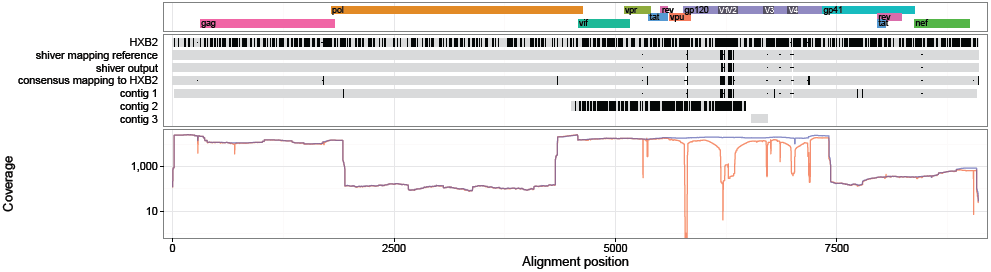
ERR732087 sequences and coverage (mapping to the `shiver` reference in blue, to HXB2 in red).

**Figure 27:**
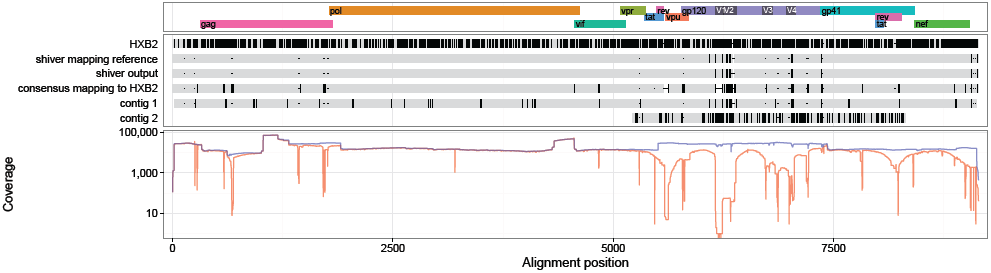
ERR732088 sequences and coverage (mapping to the `shiver` reference in blue, to HXB2 in red).

**Figure 28:**
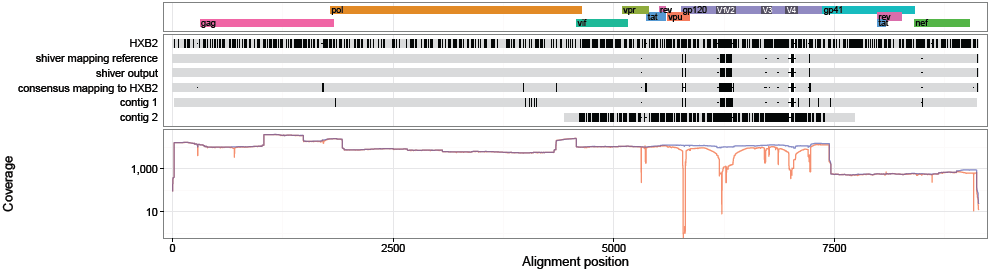
ERR732089 sequences and coverage (mapping to the `shiver` reference in blue, to HXB2 in red).

**Figure 29:**
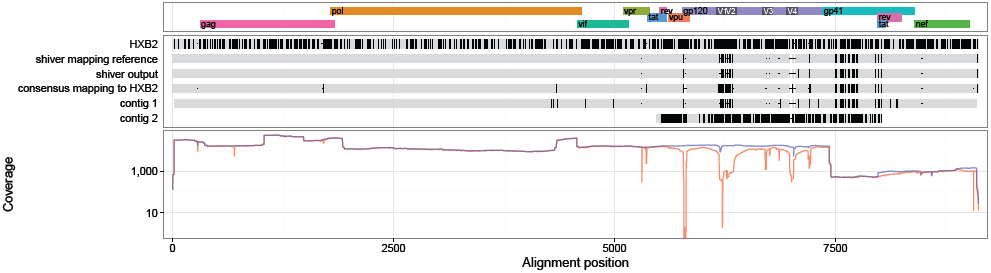
ERR732090 sequences and coverage (mapping to the `shiver` reference in blue, to HXB2 in red).

**Figure 30:**
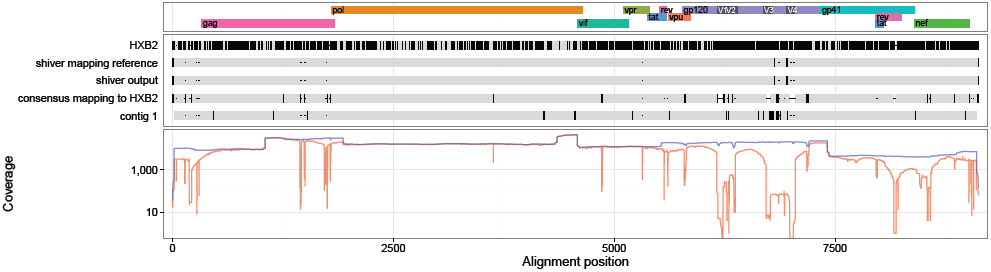
ERR732091 sequences and coverage (mapping to the `shiver` reference in blue, to HXB2 in red).

**Figure 31:**
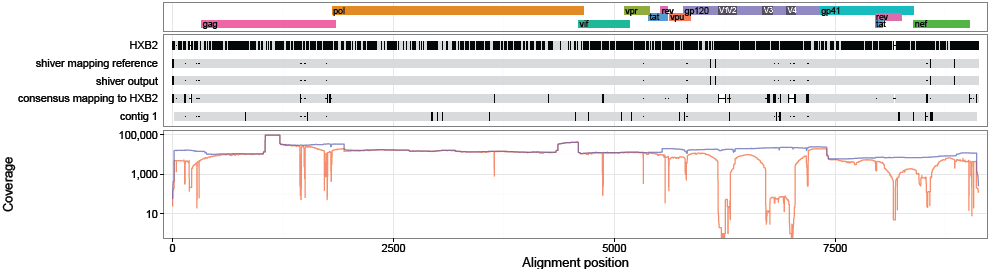
ERR732092 sequences and coverage (mapping to the `shiver` reference in blue, to HXB2 in red).

**Figure 32:**
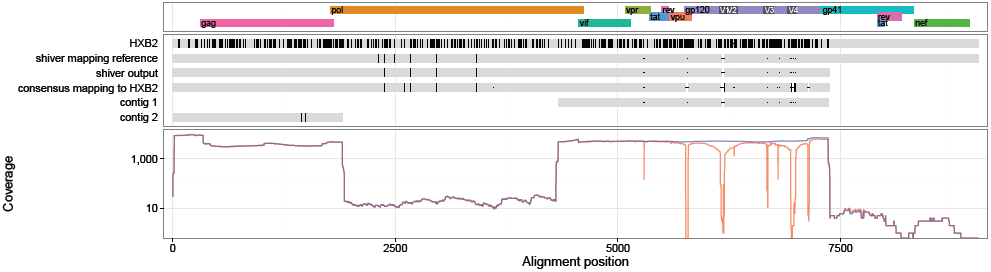
ERR732093 sequences and coverage (mapping to the `shiver` reference in blue, to HXB2 in red).

**Figure 33:**
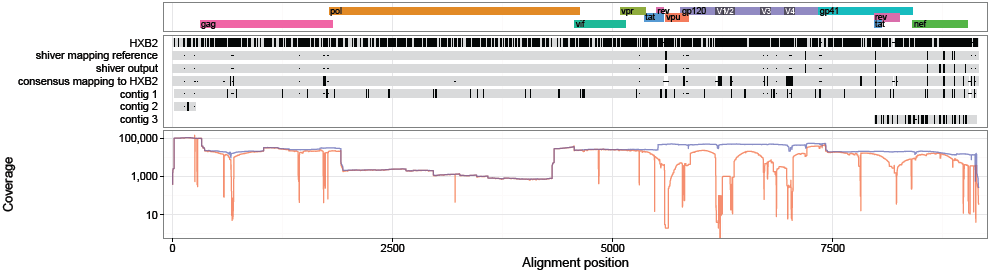
ERR732094 sequences and coverage (mapping to the `shiver` reference in blue, to HXB2 in red).

**Figure 34:**
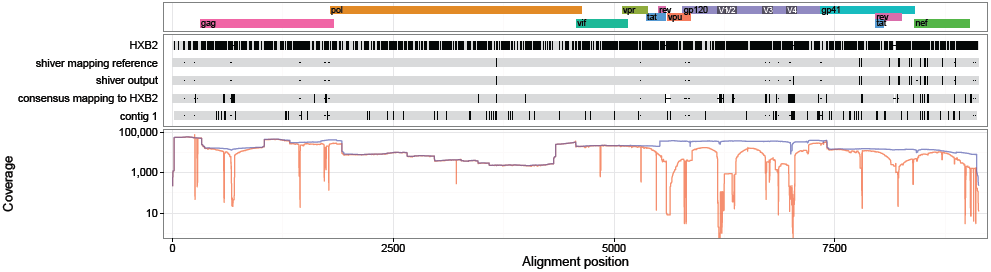
ERR732095 sequences and coverage (mapping to the `shiver` reference in blue, to HXB2 in red).

**Figure 35:**
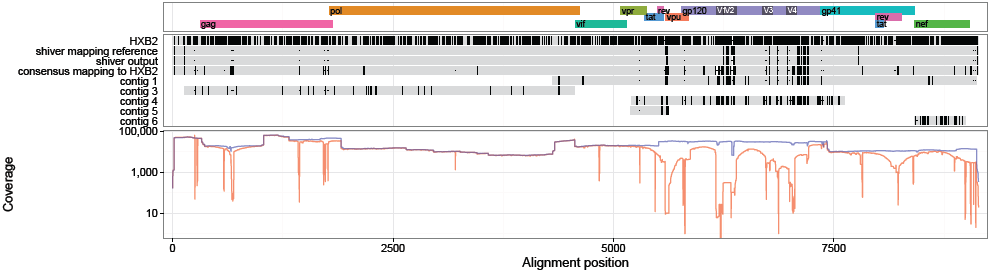
ERR732096 sequences and coverage (mapping to the `shiver` reference in blue, to HXB2 in red).

**Figure 36:**
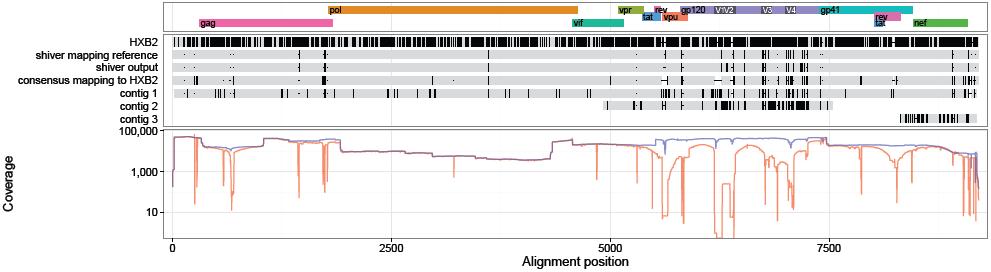
ERR732097 sequences and coverage (mapping to the `shiver` reference in blue, to HXB2 in red).

**Figure 37:**
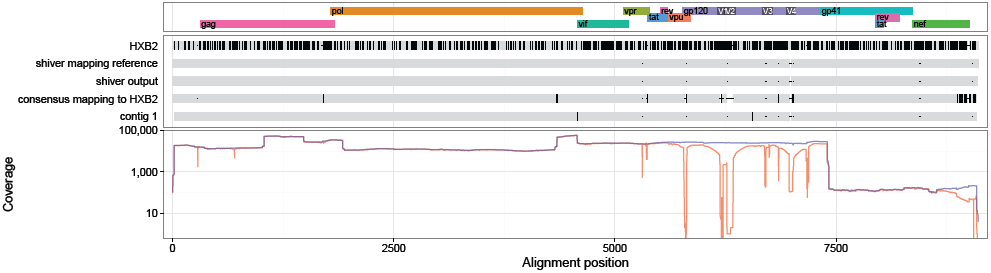
ERR732098 sequences and coverage (mapping to the `shiver` reference in blue, to HXB2 in red).

**Figure 38:**
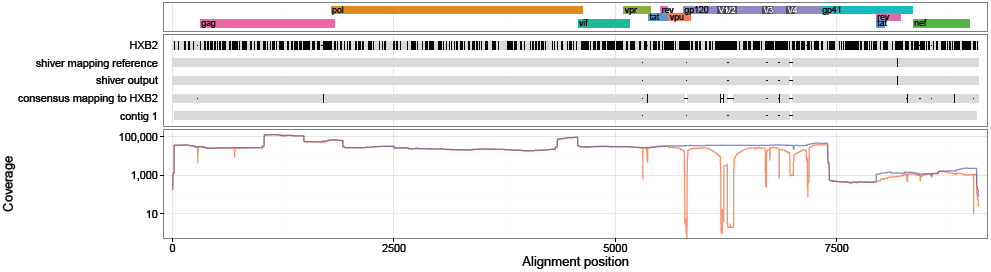
ERR732099 sequences and coverage (mapping to the `shiver` reference in blue, to HXB2 in red).

**Figure 39:**
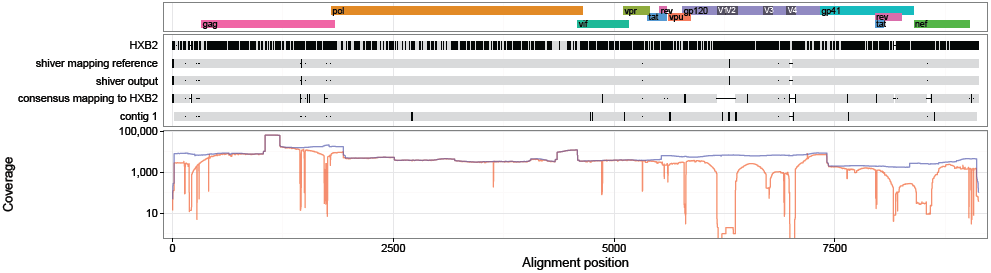
ERR732100 sequences and coverage (mapping to the `shiver` reference in blue, to HXB2 in red).

**Figure 40:**
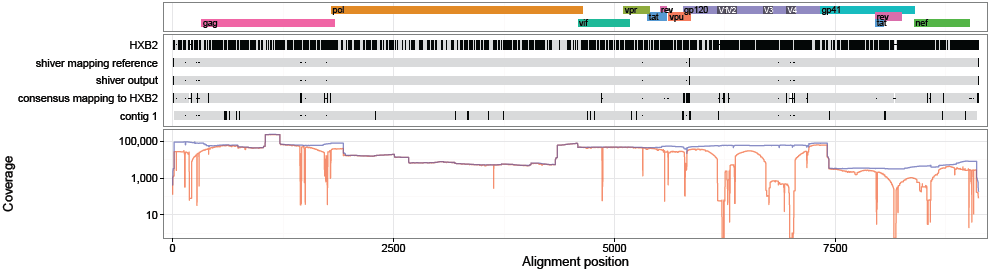
ERR732101 sequences and coverage (mapping to the `shiver` reference in blue, to HXB2 in red).

**Figure 41:**
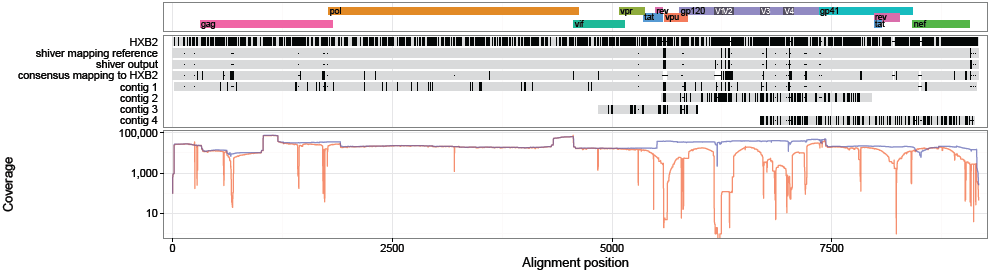
ERR732102 sequences and coverage (mapping to the `shiver` reference in blue, to HXB2 in red).

**Figure 42:**
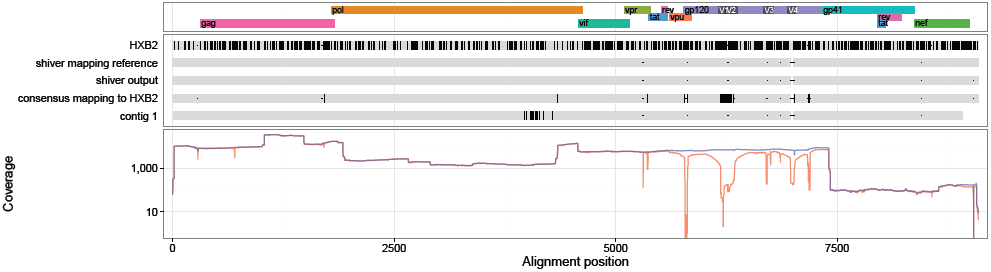
ERR732103 sequences and coverage (mapping to the `shiver` reference in blue, to HXB2 in red).

**Figure 43:**
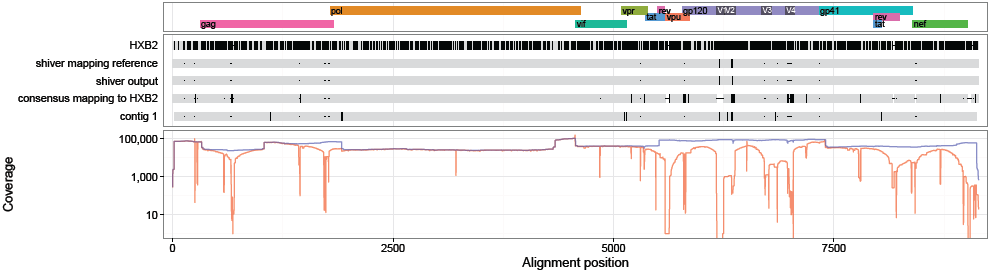
ERR732104 sequences and coverage (mapping to the `shiver` reference in blue, to HXB2 in red).

**Figure 44:**
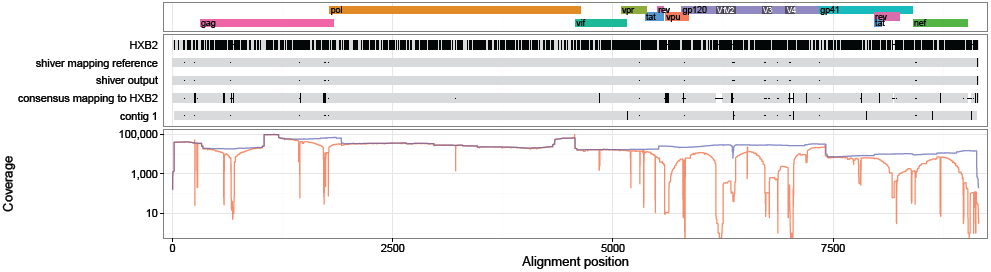
ERR732105 sequences and coverage (mapping to the `shiver` reference in blue, to HXB2 in red).

**Figure 45:**
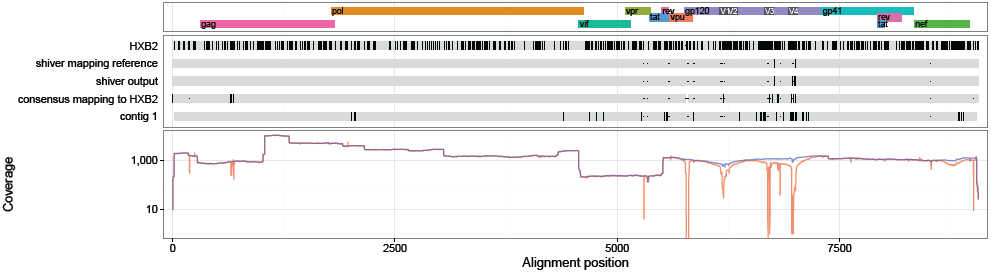
ERR732106 sequences and coverage (mapping to the `shiver` reference in blue, to HXB2 in red).

**Figure 46:**
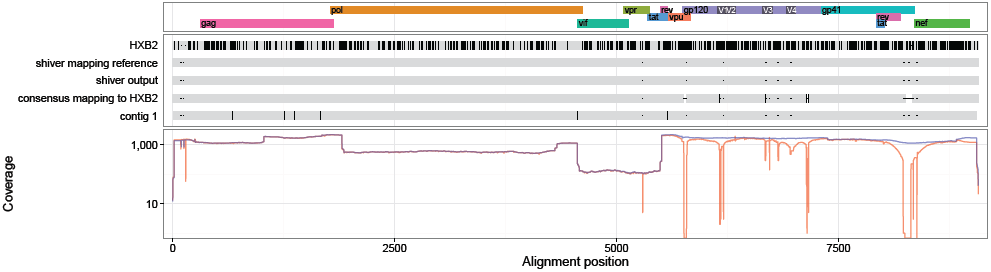
ERR732107 sequences and coverage (mapping to the `shiver` reference in blue, to HXB2 in red).

**Figure 47:**
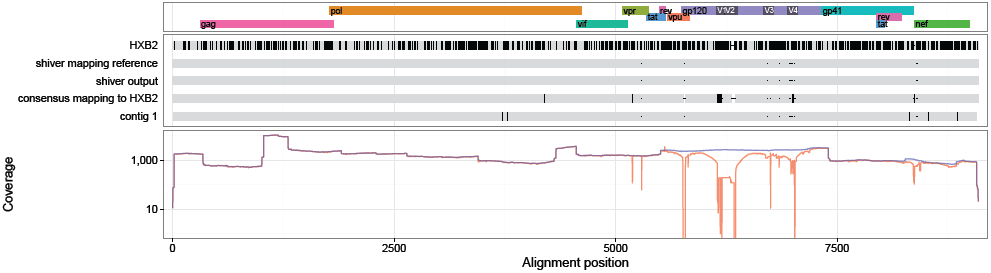
ERR732108 sequences and coverage (mapping to the `shiver` reference in blue, to HXB2 in red).

**Figure 48:**
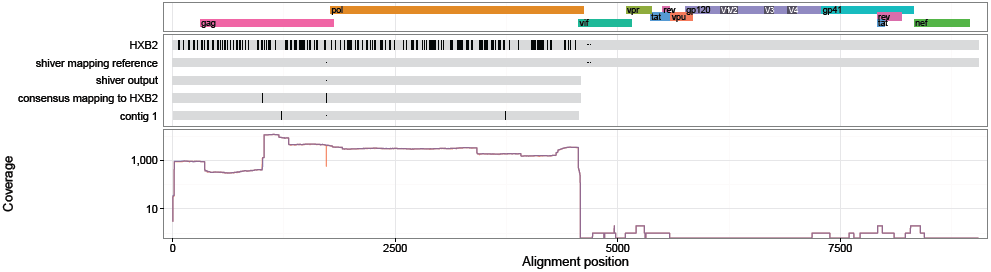
ERR732109 sequences and coverage (mapping to the `shiver` reference in blue, to HXB2 in red).

**Figure 49:**
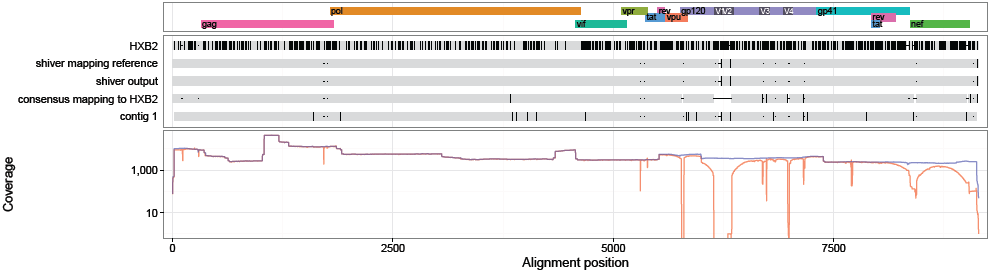
ERR732110 sequences and coverage (mapping to the `shiver` reference in blue, to HXB2 in red).

**Figure 50:**
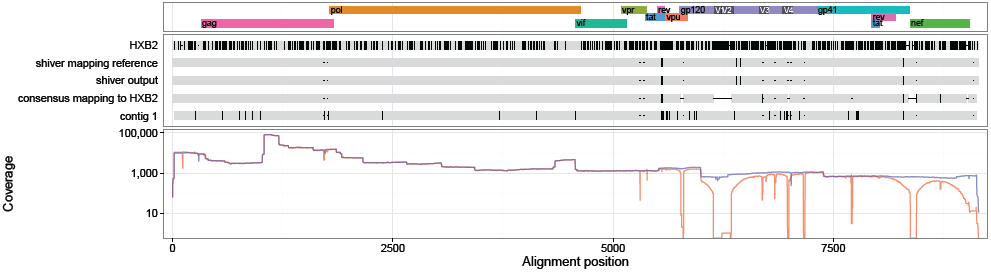
ERR732111 sequences and coverage (mapping to the `shiver` reference in blue, to HXB2 in red).

**Figure 51:**
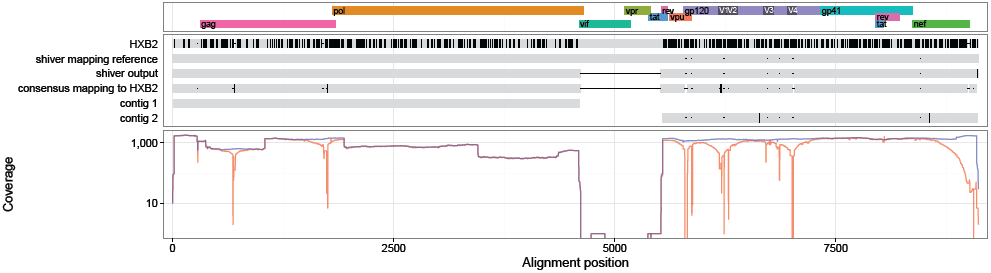
ERR732112 sequences and coverage (mapping to the `shiver` reference in blue, to HXB2 in red).

**Figure 52:**
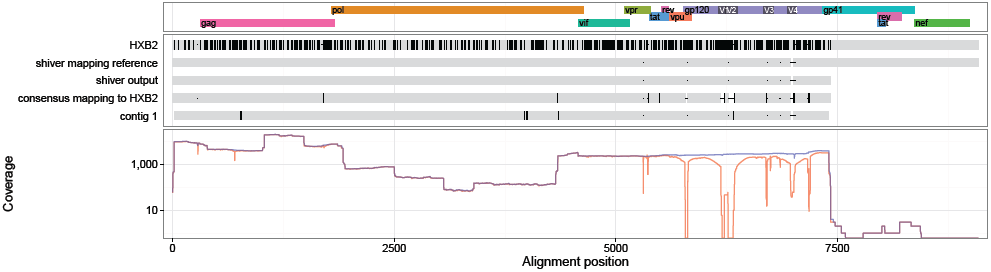
ERR732113 sequences and coverage (mapping to the `shiver` reference in blue, to HXB2 in red).

**Figure 53:**
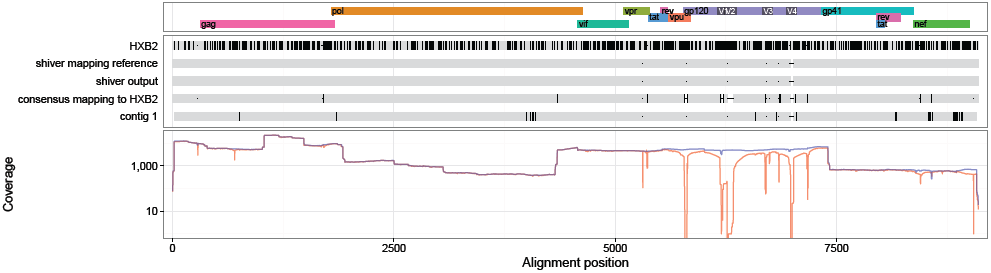
ERR732114 sequences and coverage (mapping to the `shiver` reference in blue, to HXB2 in red).

**Figure 54:**
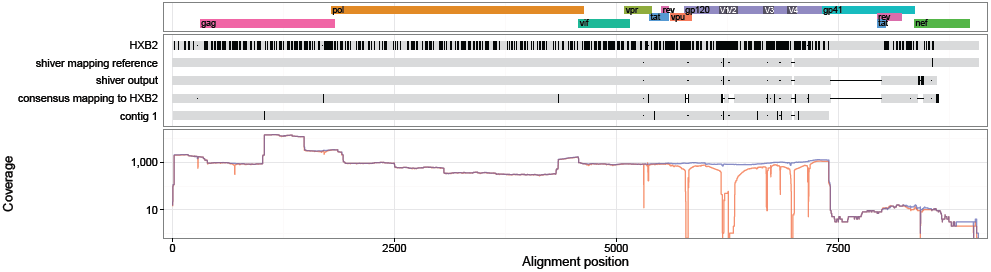
ERR732115 sequences and coverage (mapping to the `shiver` reference in blue, to HXB2 in red).

**Figure 55:**
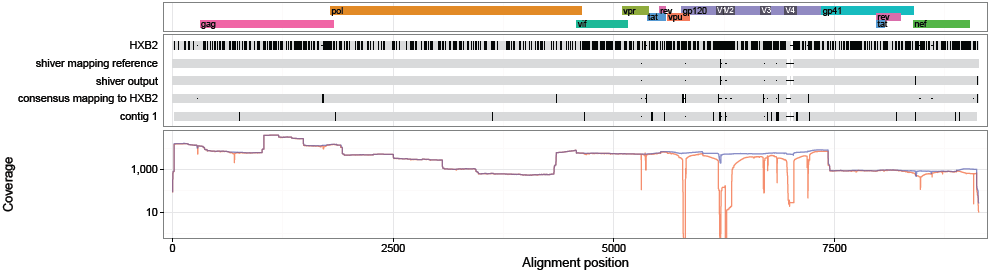
ERR732116 sequences and coverage (mapping to the `shiver` reference in blue, to HXB2 in red).

**Figure 56:**
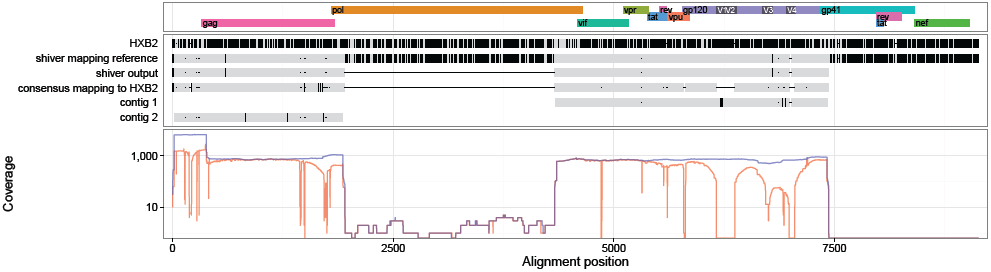
ERR732117 sequences and coverage (mapping to the `shiver` reference in blue, to HXB2 in red).

**Figure 57:**
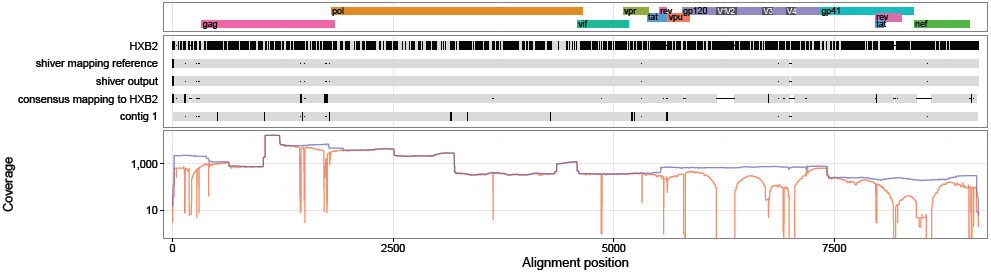
ERR732118 sequences and coverage (mapping to the `shiver` reference in blue, to HXB2 in red).

**Figure 58:**
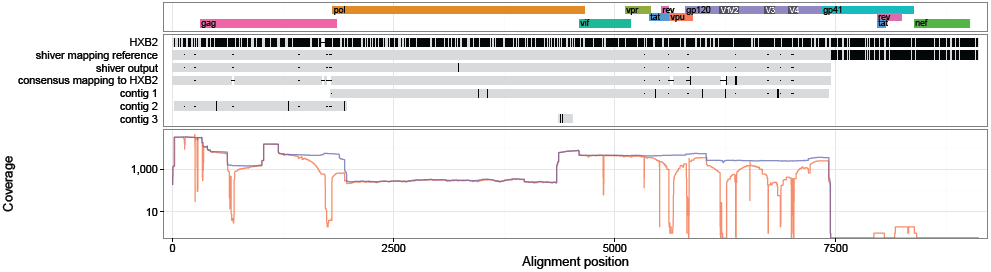
ERR732119 sequences and coverage (mapping to the `shiver` reference in blue, to HXB2 in red).

**Figure 59:**
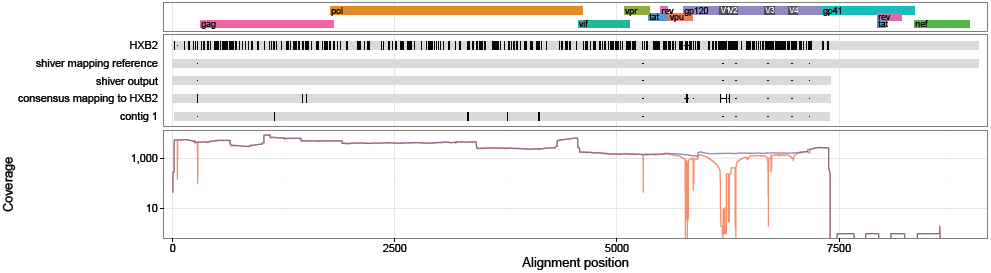
ERR732120 sequences and coverage (mapping to the `shiver` reference in blue, to HXB2 in red).

**Figure 60:**
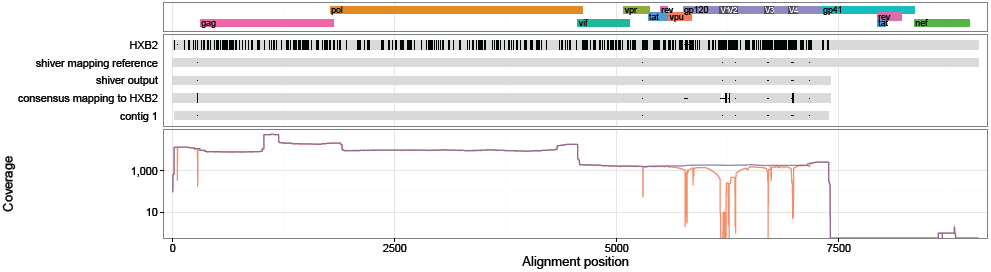
ERR732121 sequences and coverage (mapping to the `shiver` reference in blue, to HXB2 in red).

**Figure 61:**
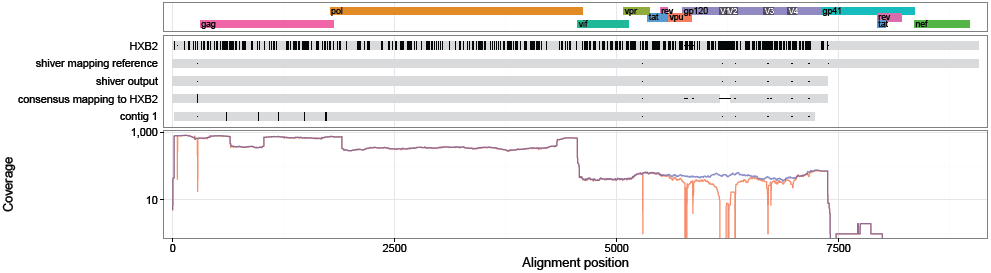
ERR732122 sequences and coverage (mapping to the `shiver` reference in blue, to HXB2 in red).

**Figure 62:**
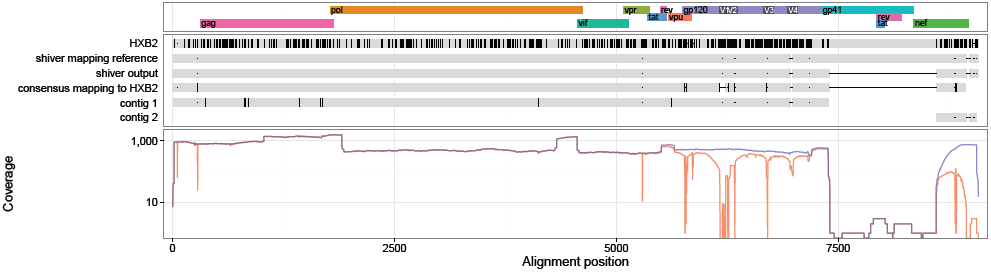
ERR732123 sequences and coverage (mapping to the `shiver` reference in blue, to HXB2 in red).

**Figure 63:**
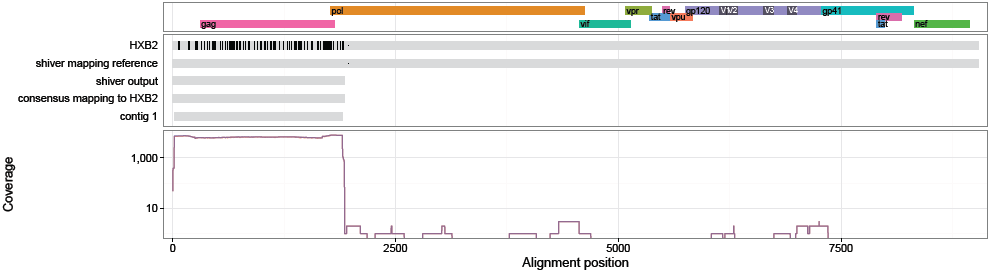
ERR732124 sequences and coverage (mapping to the `shiver` reference in blue, to HXB2 in red).

**Figure 64:**
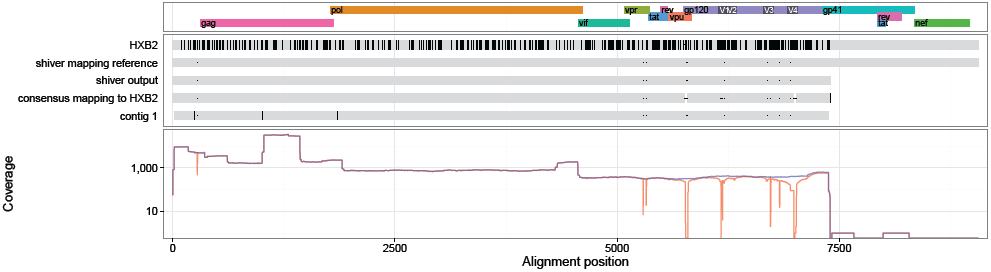
ERR732126 sequences and coverage (mapping to the `shiver` reference in blue, to HXB2 in red).

**Figure 65:**
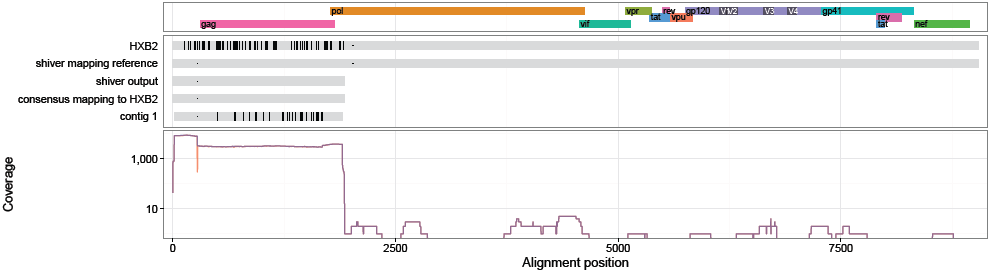
ERR732127 sequences and coverage (mapping to the `shiver` reference in blue, to HXB2 in red).

**Figure 66:**
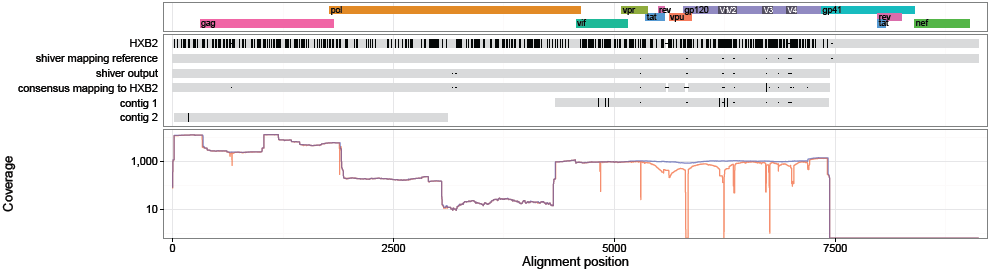
ERR732128 sequences and coverage (mapping to the `shiver` reference in blue, to HXB2 in red).

**Figure 67:**
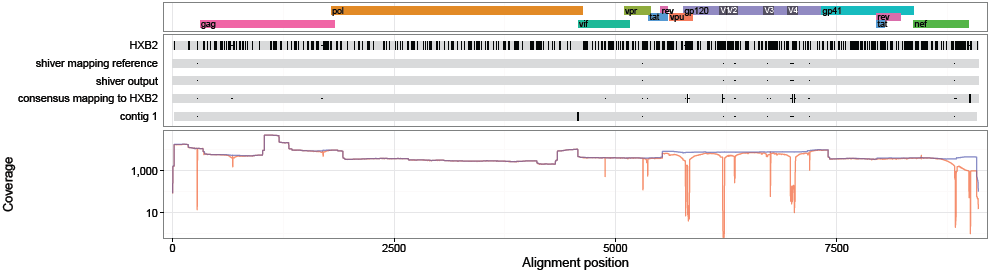
ERR732129 sequences and coverage (mapping to the `shiver` reference in blue, to HXB2 in red).

**Figure 68:**
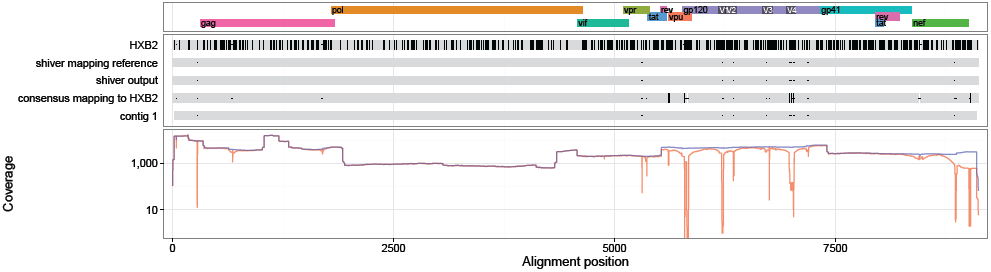
ERR732130 sequences and coverage (mapping to the `shiver` reference in blue, to HXB2 in red).

**Figure 69:**
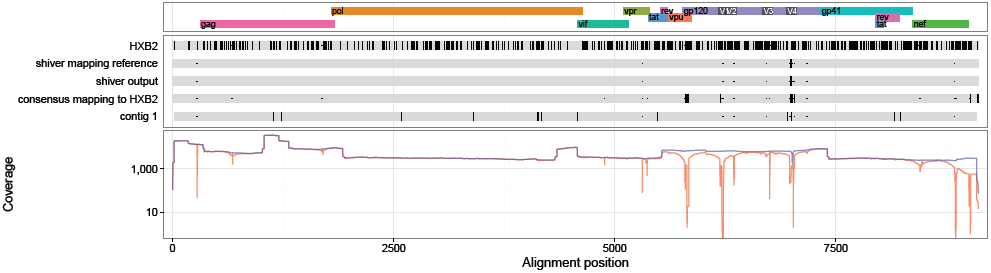
ERR732131 sequences and coverage (mapping to the `shiver` reference in blue, to HXB2 in red).

**Figure 70:**
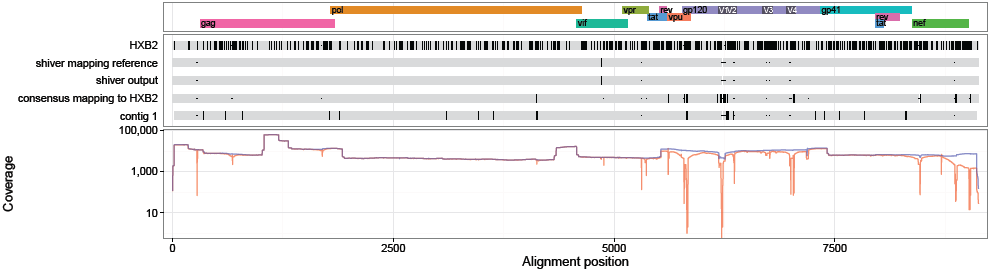
ERR732132 sequences and coverage (mapping to the `shiver` reference in blue, to HXB2 in red).

## Appendix G Sequences and Coverage by Sample: Hiseq Data

Plots of the same format as those described in Appendix F.

**Figure 71:**
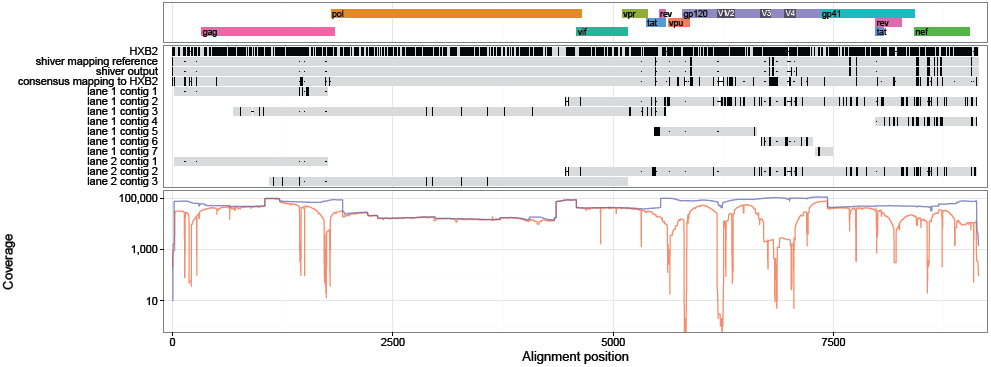
17621_3_80 sequences and coverage (mapping to the `shiver` reference in blue, to HXB2 in red).

**Figure 72:**
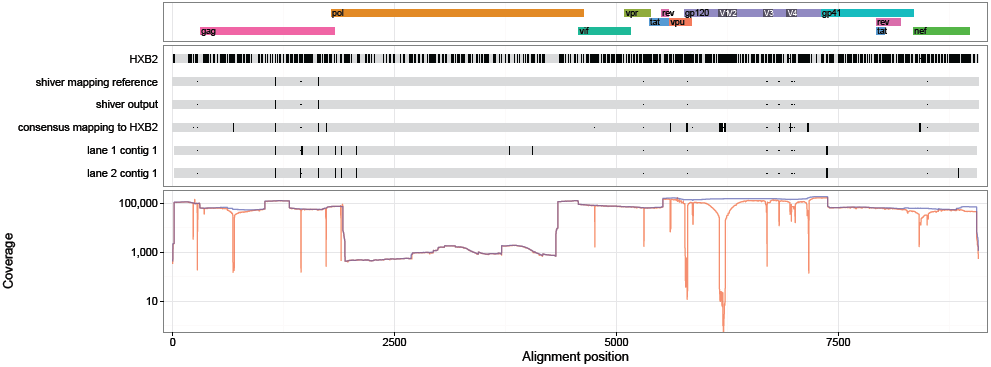
17653_3_25 sequences and coverage (mapping to the `shiver` reference in blue, to HXB2 in red).

**Figure 73:**
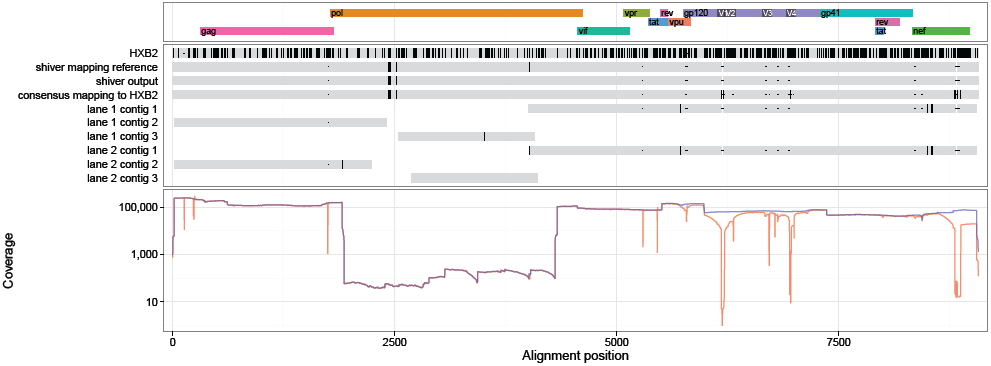
17653 _3_ 36 sequences and coverage (mapping to the `shiver` reference in blue, to HXB2 in red).

**Figure 74:**
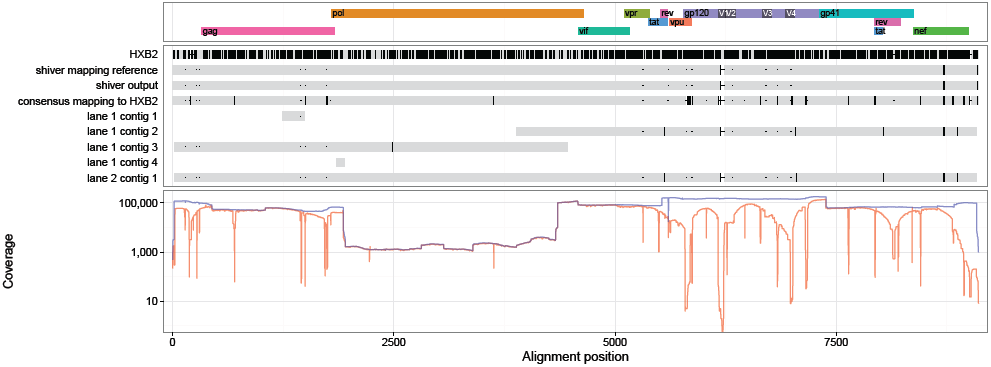
17653_3_56 sequences and coverage (mapping to the `shiver` reference in blue, to HXB2 in red).

**Figure 75:**
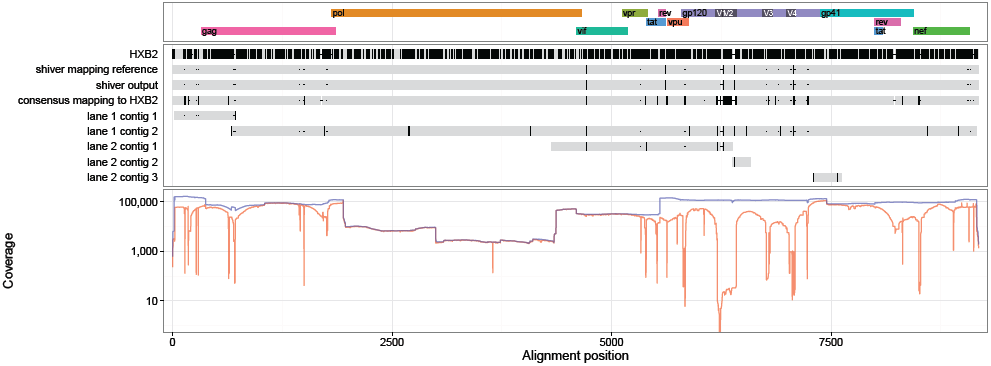
17653_3_62 sequences and coverage (mapping to the `shiver` reference in blue, to HXB2 in red).

**Figure 76:**
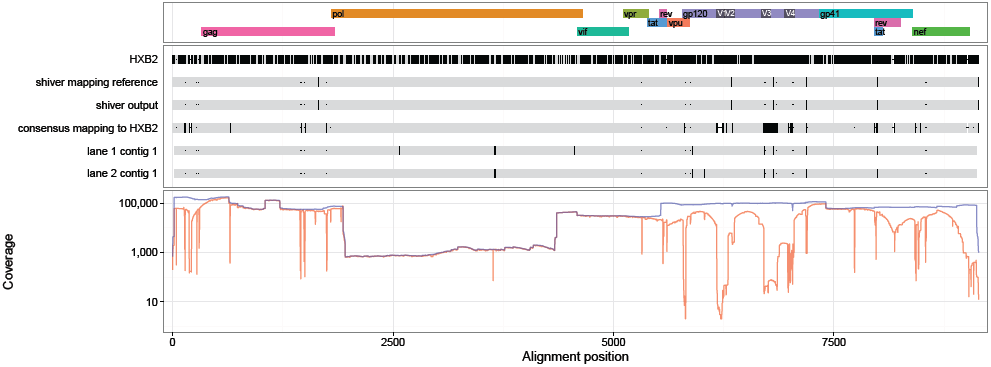
17653_3_64 sequences and coverage (mapping to the `shiver` reference in blue, to HXB2 in red).

**Figure 77:**
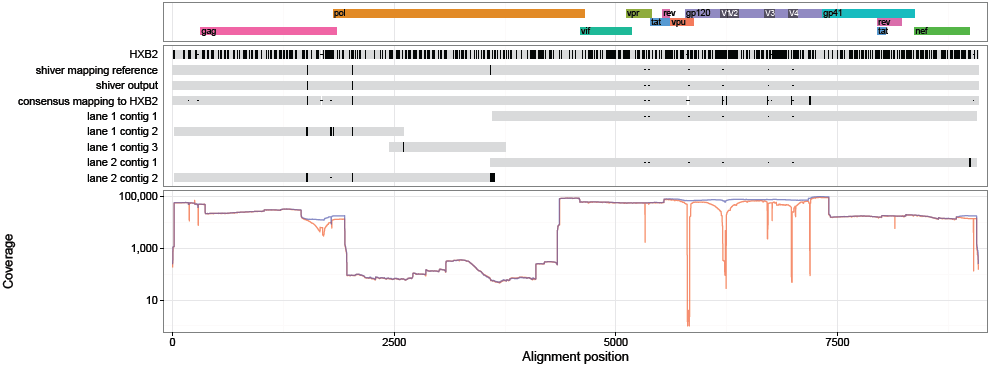
17653_3_72 sequences and coverage (mapping to the `shiver` reference in blue, to HXB2 in red).

**Figure 78:**
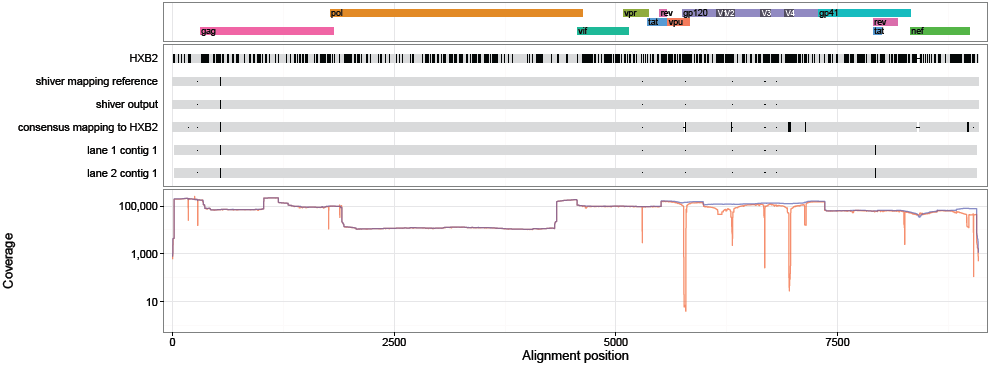
17653_3_74 sequences and coverage (mapping to the `shiver` reference in blue, to HXB2 in red).

**Figure 79:**
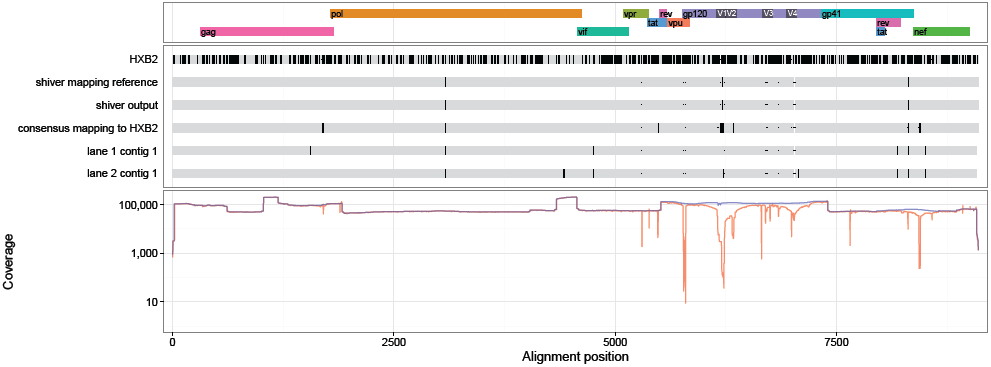
17654_3_46 sequences and coverage (mapping to the `shiver` reference in blue, to HXB2 in red).

**Figure 80:**
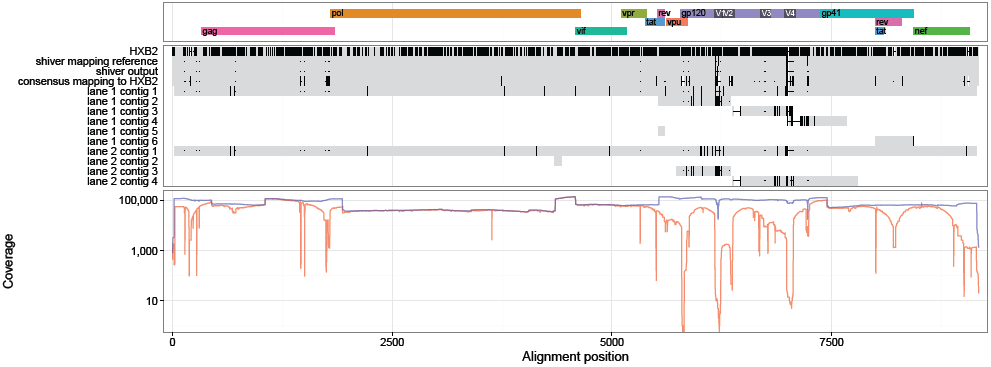
17654_3_71 sequences and coverage (mapping to the `shiver` reference in blue, to HXB2 in red).

**Figure 81:**
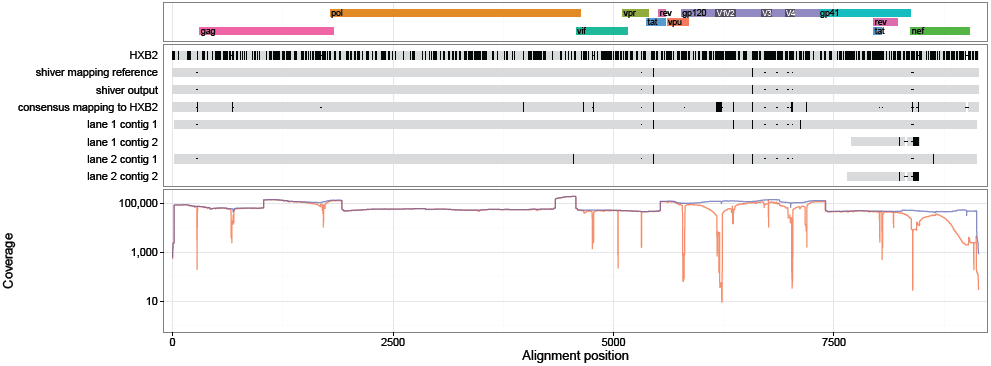
17654_3_72 sequences and coverage (mapping to the `shiver` reference in blue, to HXB2 in red).

**Figure 82:**
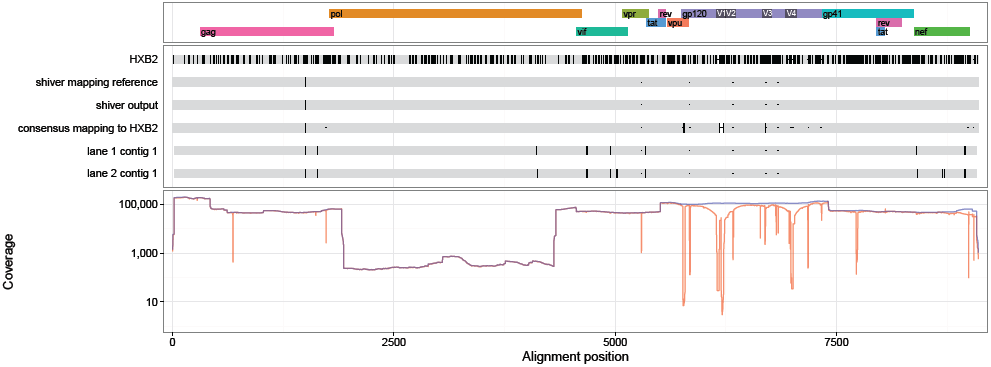
17654_3_78 sequences and coverage (mapping to the `shiver` reference in blue, to HXB2 in red).

**Figure 83:**
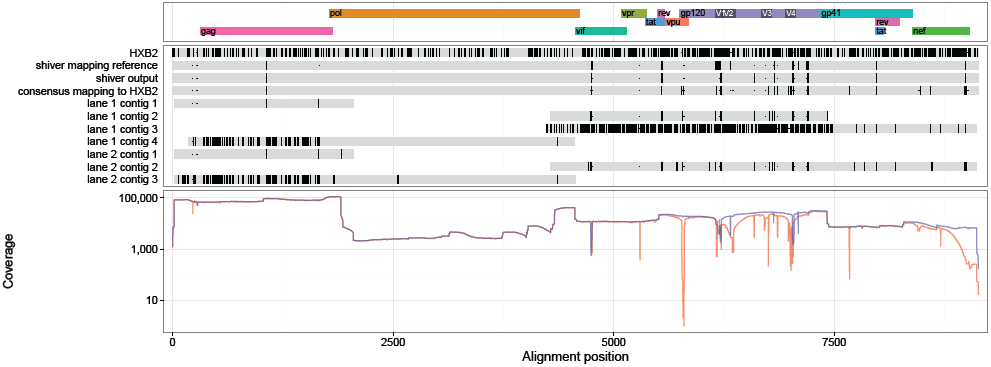
17795_3_40 sequences and coverage (mapping to the `shiver` reference in blue, to HXB2 in red).

**Figure 84:**
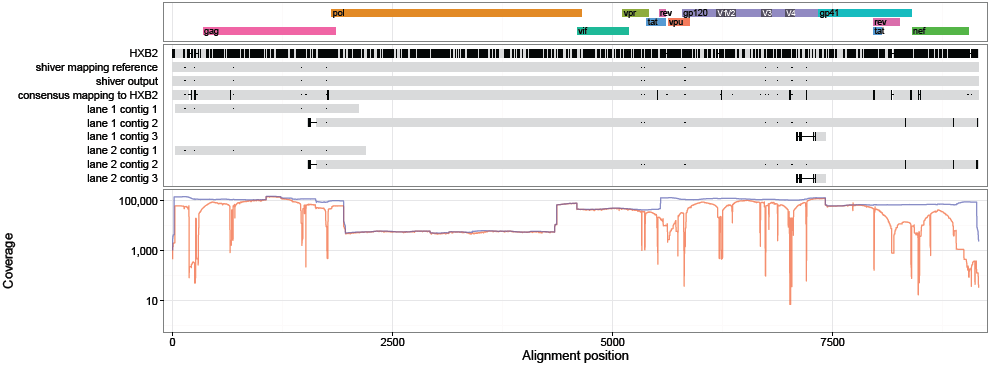
17796_3_1 sequences and coverage (mapping to the `shiver` reference in blue, to HXB2 in red).

**Figure 85:**
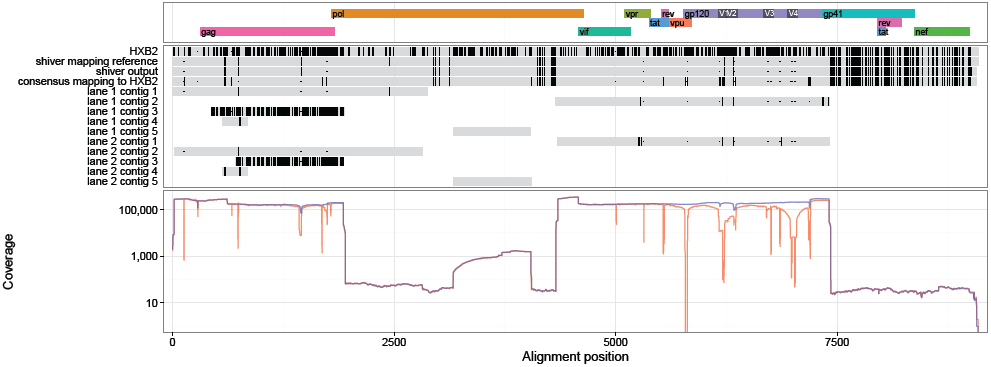
17796_3_29 sequences and coverage (mapping to the `shiver` reference in blue, to HXB2 in red).

**Figure 86:**
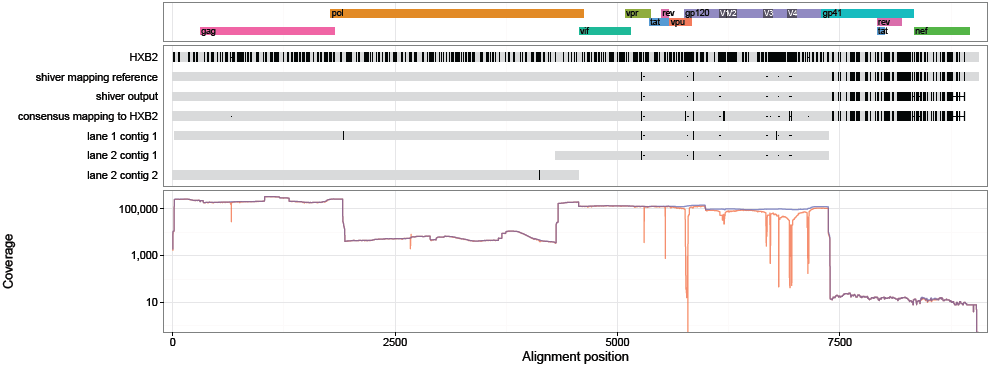
17796_3_30 sequences and coverage (mapping to the `shiver` reference in blue, to HXB2 in red).

**Figure 87:**
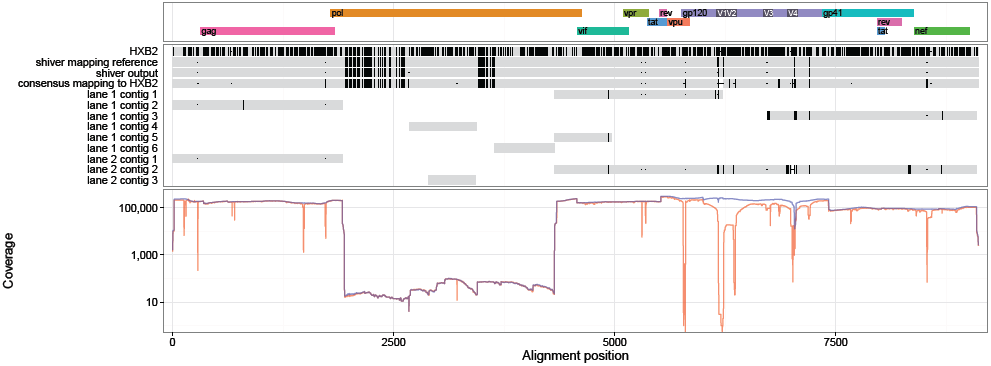
17796_3_35 sequences and coverage (mapping to the `shiver` reference in blue, to HXB2 in red).

**Figure 88:**
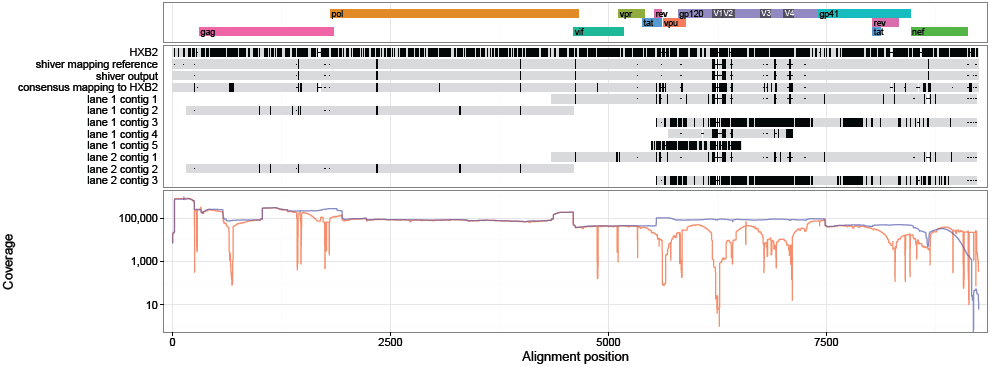
18209_3_31 sequences and coverage (mapping to the `shiver` reference in blue, to HXB2 in red).

**Figure 89:**
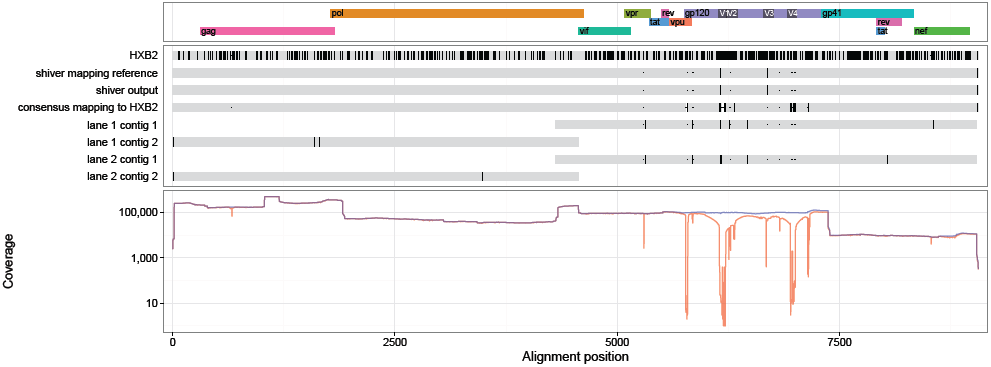
18209_3_36 sequences and coverage (mapping to the `shiver` reference in blue, to HXB2 in red).

**Figure 90:**
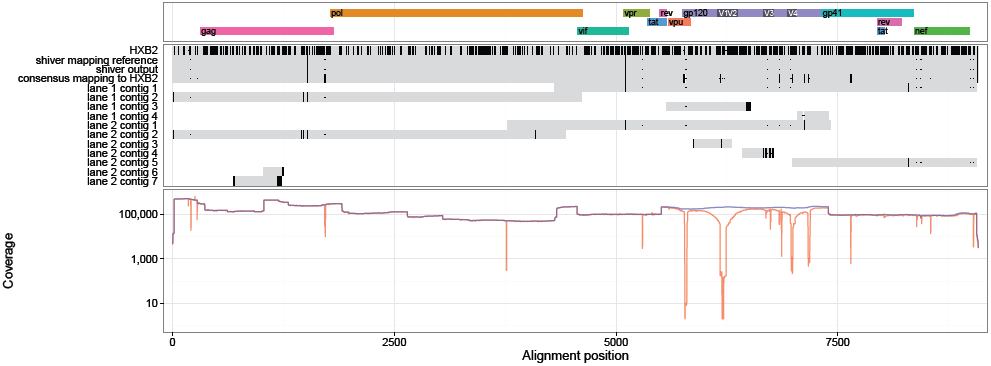
18209_3_38 sequences and coverage (mapping to the `shiver` reference in blue, to HXB2 in red).

**Figure 91:**
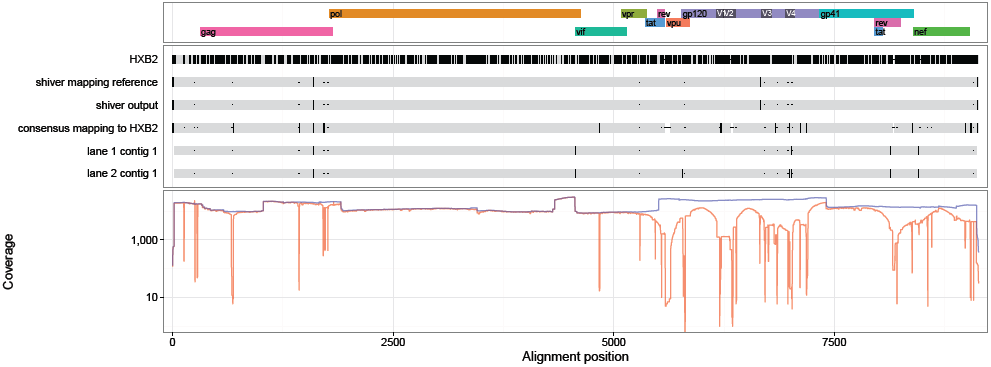
19561_3_127 sequences and coverage (mapping to the `shiver` reference in blue, to HXB2 in red).

**Figure 92:**
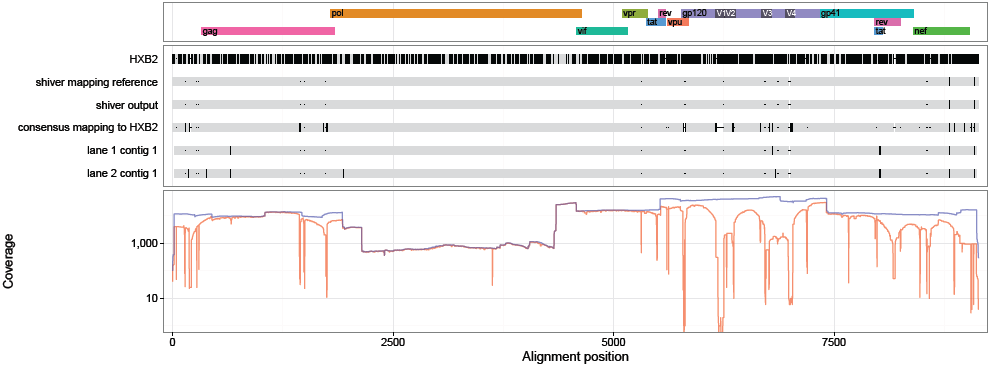
19562_3_109 sequences and coverage (mapping to the `shiver` reference in blue, to HXB2 in red).

**Figure 93:**
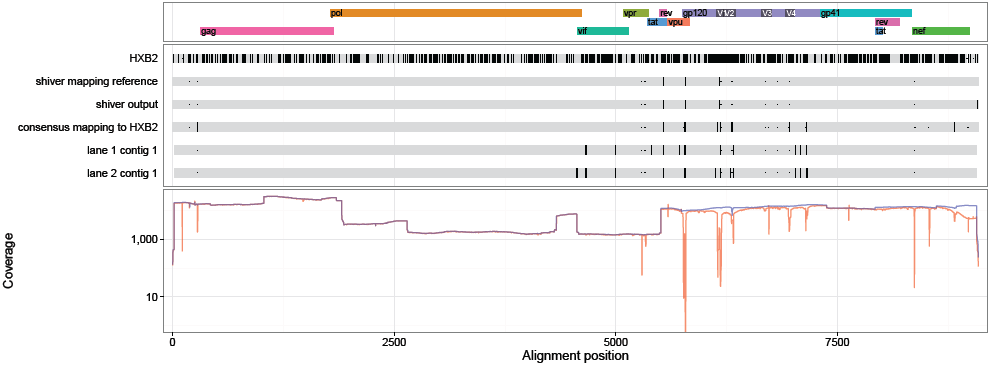
19562_3_2 sequences and coverage (mapping to the `shiver` reference in blue, to HXB2 in red).

**Figure 94:**
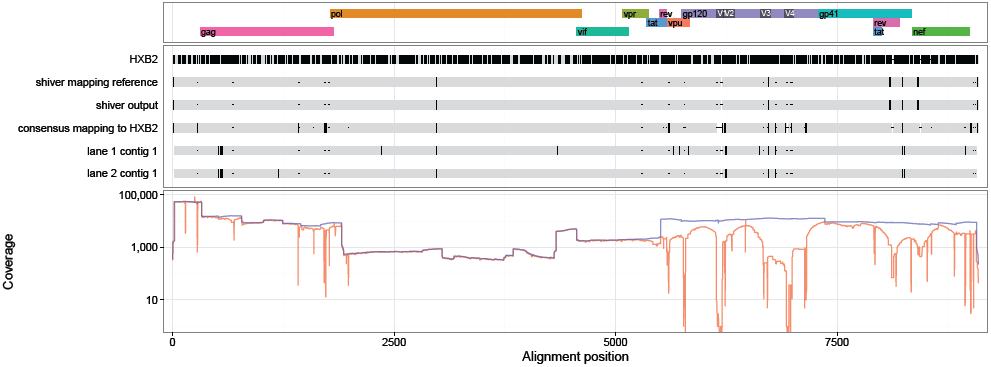
19562_3_30 sequences and coverage (mapping to the `shiver` reference in blue, to HXB2 in red).

**Figure 95:**
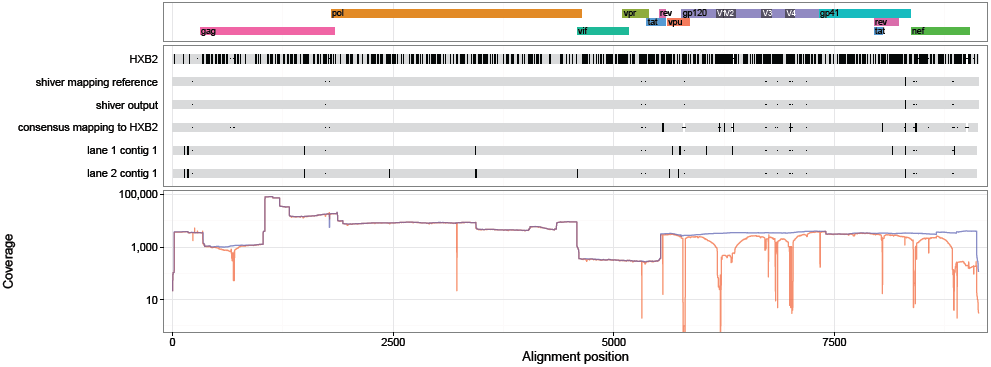
19562_3_31 sequences and coverage (mapping to the `shiver` reference in blue, to HXB2 in red).

**Figure 96:**
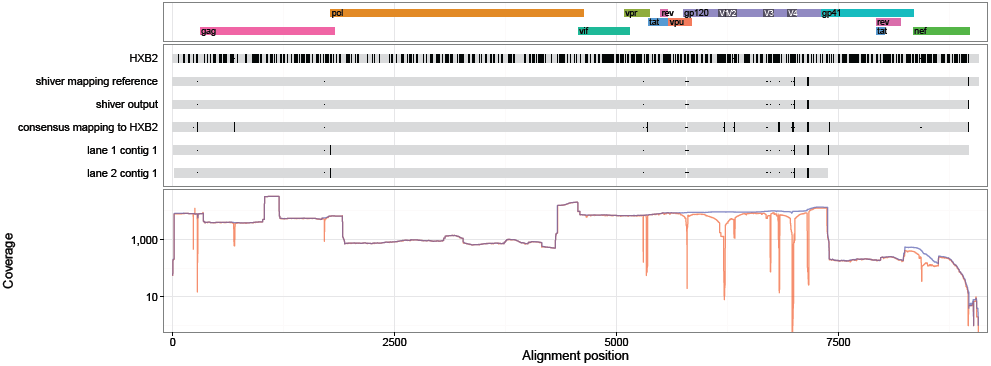
19562_3_46 sequences and coverage (mapping to the `shiver` reference in blue, to HXB2 in red).

**Figure 97:**
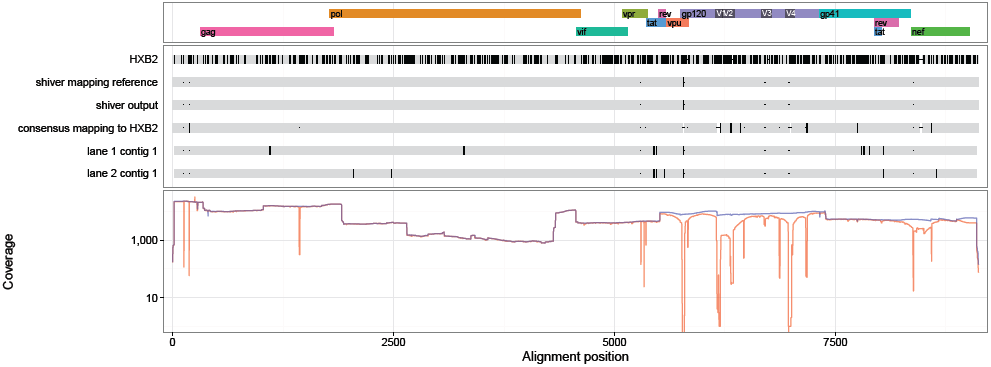
19562_3_50 sequences and coverage (mapping to the `shiver` reference in blue, to HXB2 in red).

**Figure 98:**
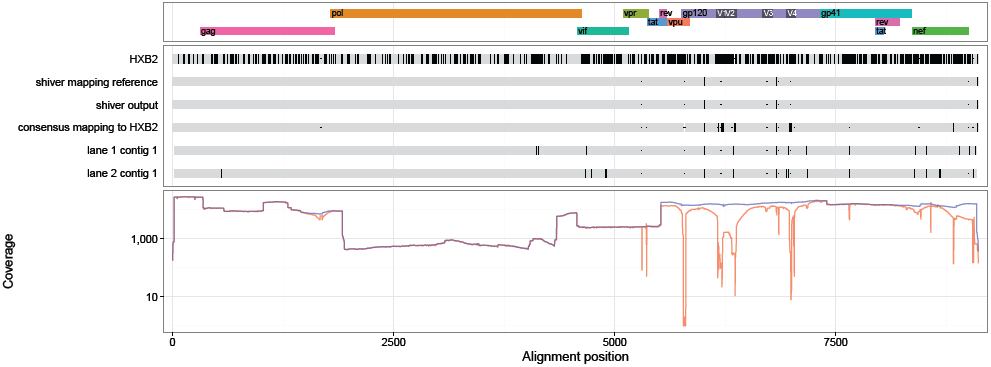
19562_3_51 sequences and coverage (mapping to the `shiver` reference in blue, to HXB2 in red).

**Figure 99:**
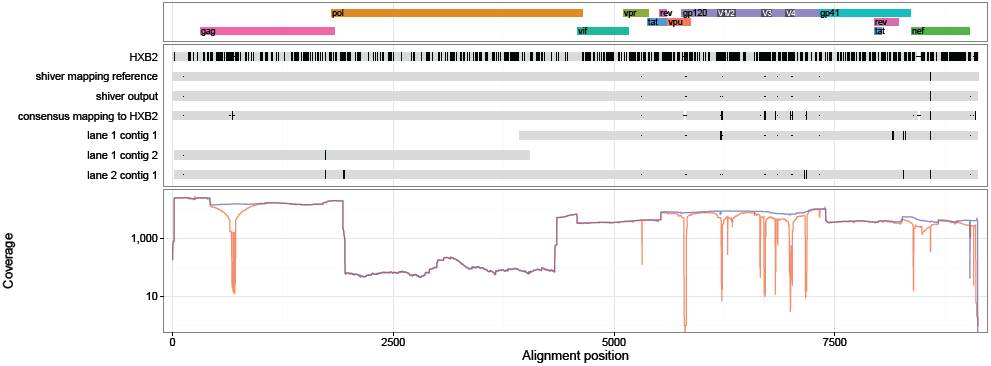
19562_3_6 sequences and coverage (mapping to the `shiver` reference in blue, to HXB2 in red).

**Figure 100:**
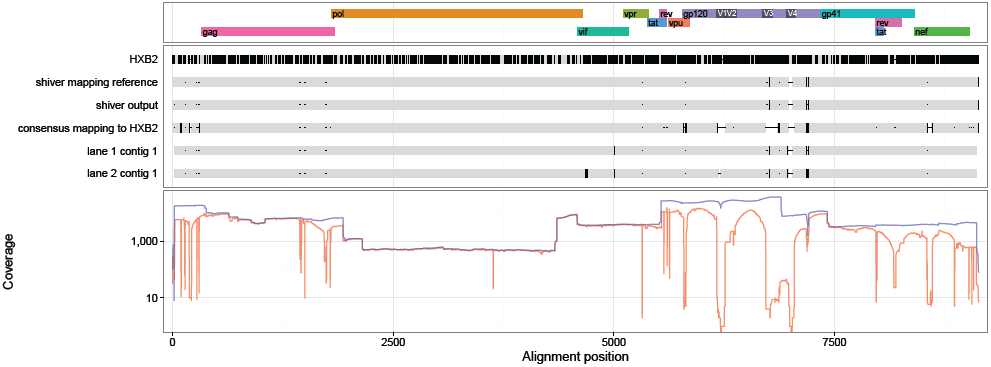
19893_3_71 sequences and coverage (mapping to the `shiver` reference in blue, to HXB2 in red).

**Figure 101:**
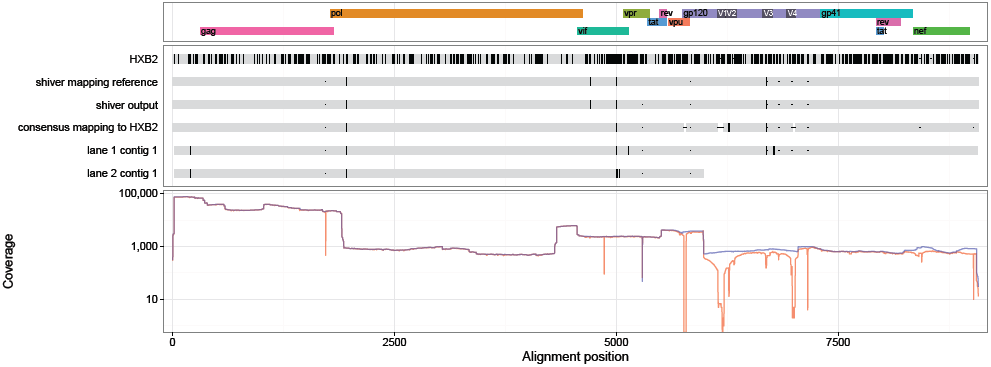
19960_3_116 sequences and coverage (mapping to the `shiver` reference in blue, to HXB2 in red).

**Figure 102:**
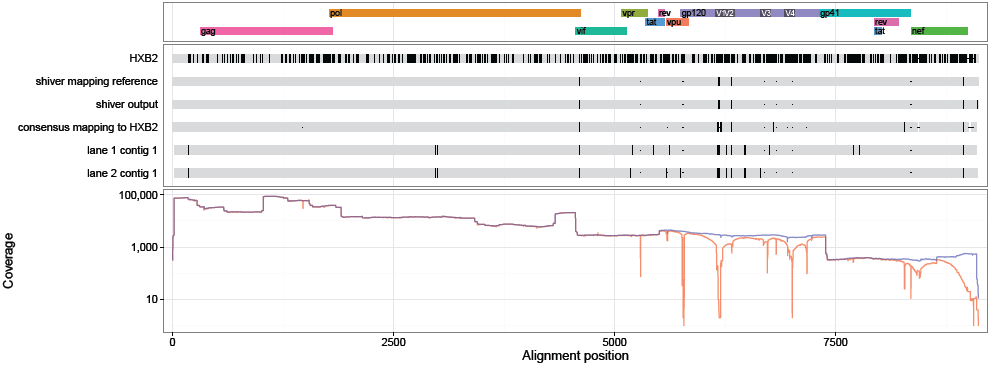
19960_3_119 sequences and coverage (mapping to the `shiver` reference in blue, to HXB2 in red).

**Figure 103:**
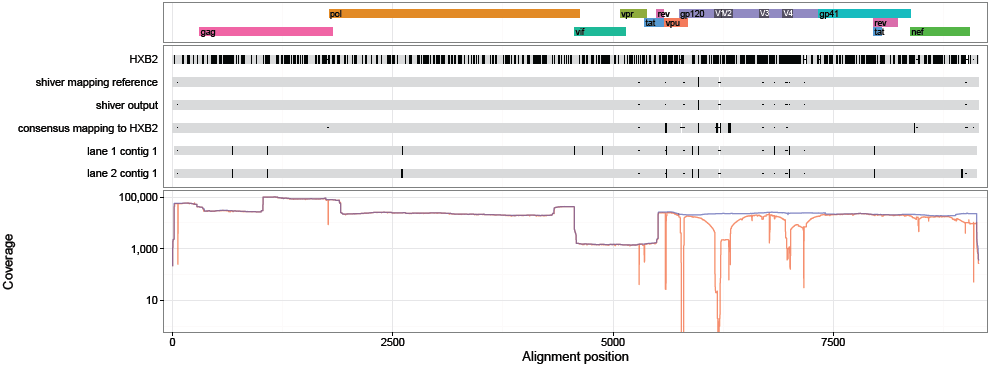
19960_3_11 sequences and coverage (mapping to the `shiver` reference in blue, to HXB2 in red).

**Figure 104:**
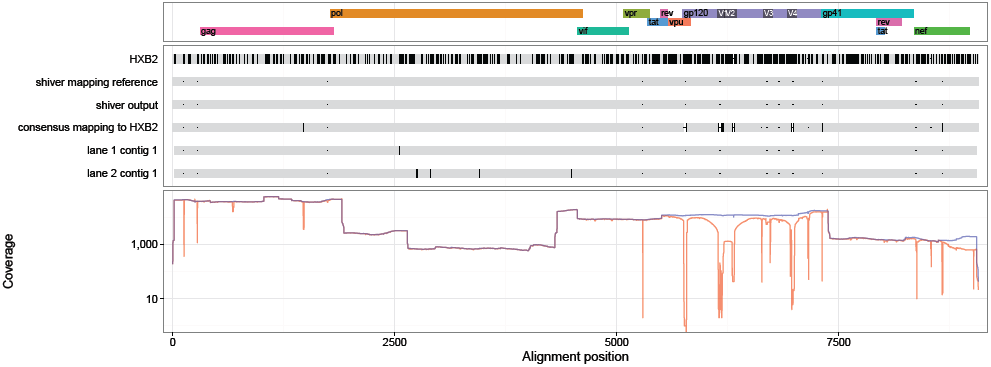
19960_3_12 sequences and coverage (mapping to the `shiver` reference in blue, to HXB2 in red).

**Figure 105:**
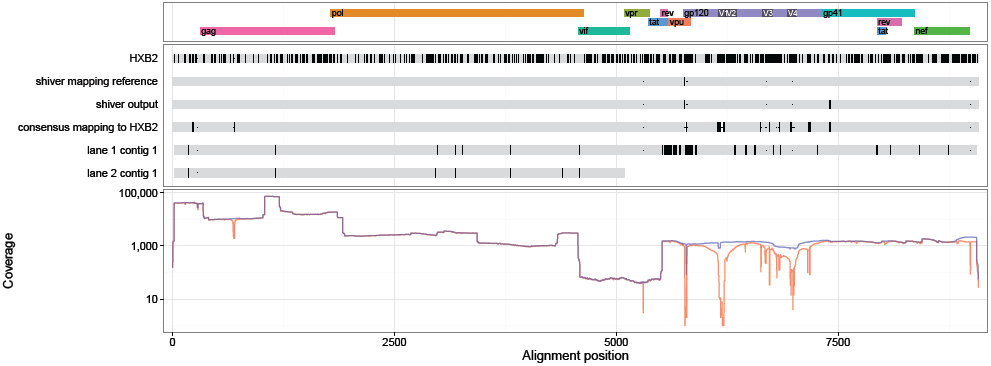
19960_3_146 sequences and coverage (mapping to the `shiver` reference in blue, to HXB2 in red).

**Figure 106:**
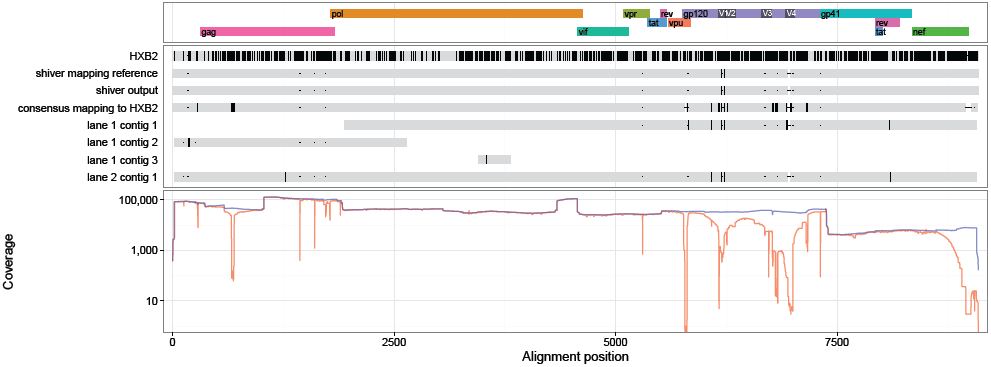
19960_3_15 sequences and coverage (mapping to the `shiver` reference in blue, to HXB2 in red).

**Figure 107:**
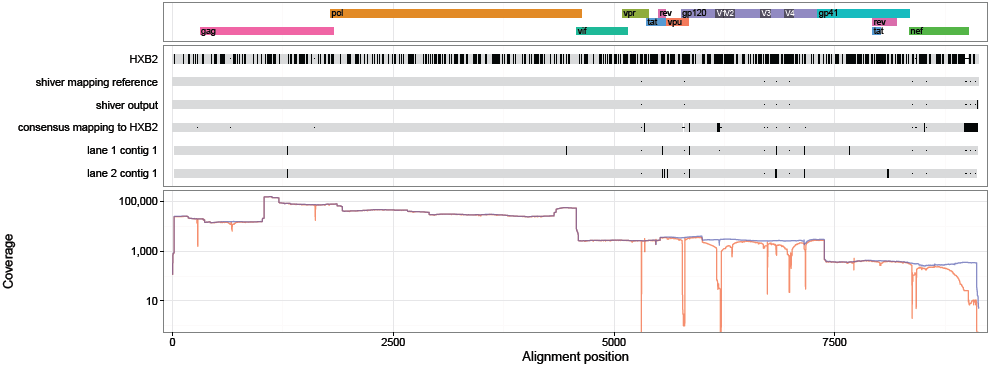
19960_3_16 sequences and coverage (mapping to the `shiver` reference in blue, to HXB2 in red).

**Figure 108:**
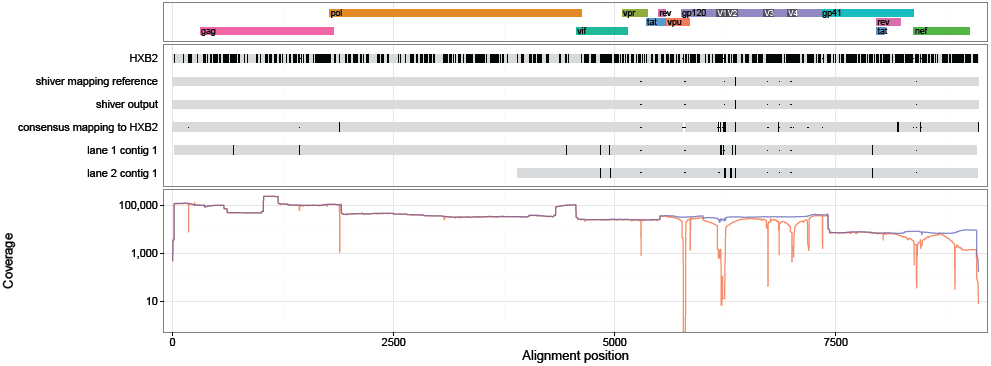
19960_3_17 sequences and coverage (mapping to the `shiver` reference in blue, to HXB2 in red).

**Figure 109:**
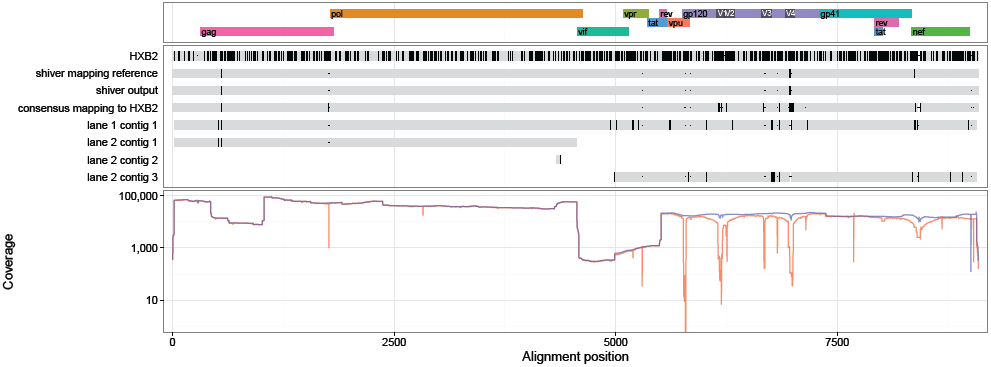
19960_3_18 sequences and coverage (mapping to the `shiver` reference in blue, to HXB2 in red).

**Figure 110:**
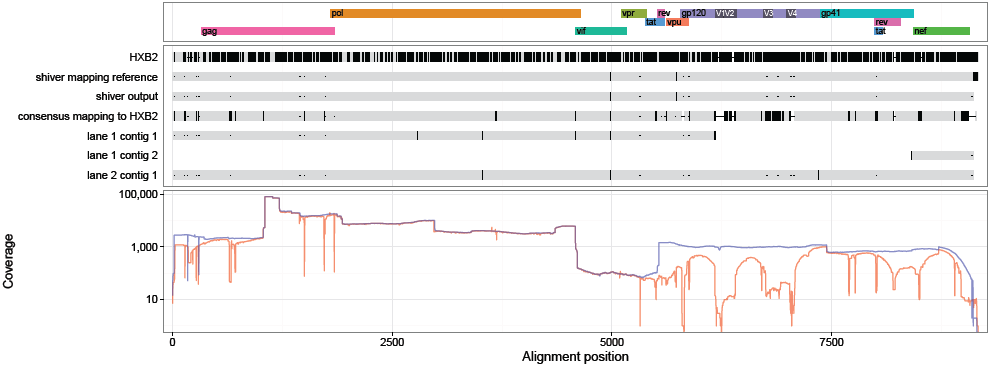
19960_3_22 sequences and coverage (mapping to the `shiver` reference in blue, to HXB2 in red).

**Figure 111:**
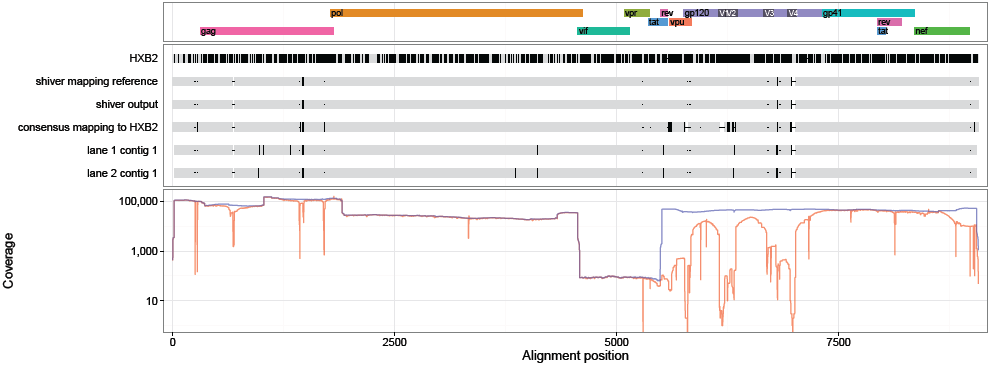
19960_3_28 sequences and coverage (mapping to the `shiver` reference in blue, to HXB2 in red).

**Figure 112:**
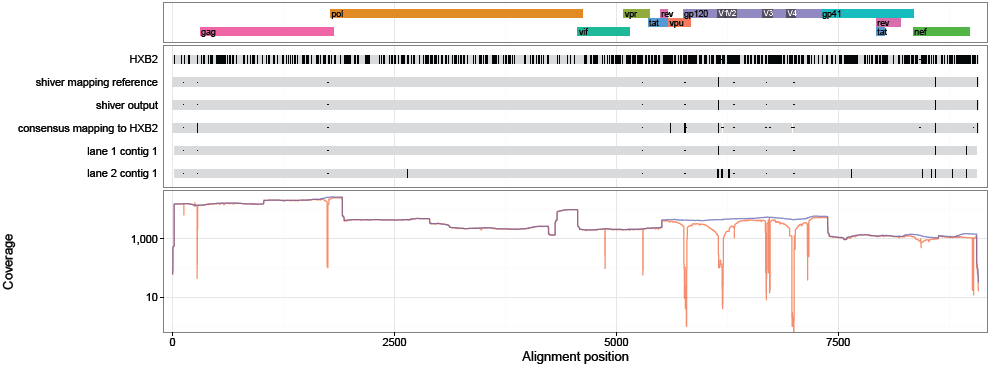
19960_3_40 sequences and coverage (mapping to the `shiver` reference in blue, to HXB2 in red).

**Figure 113:**
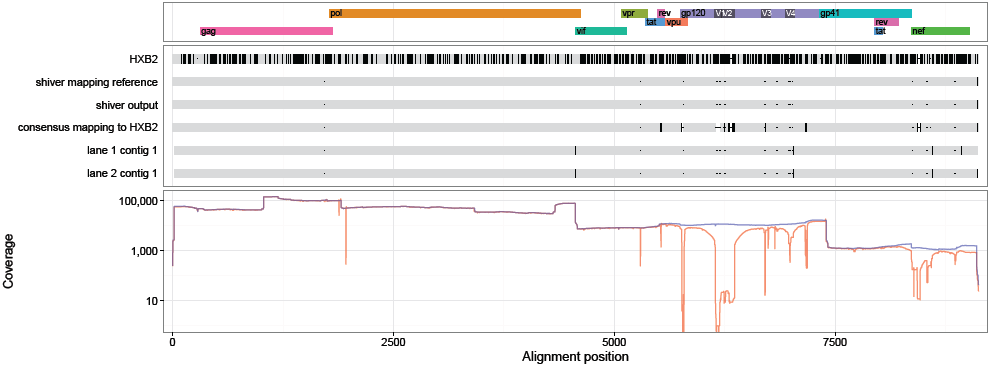
19960_3_44 sequences and coverage (mapping to the `shiver` reference in blue, to HXB2 in red).

**Figure 114:**
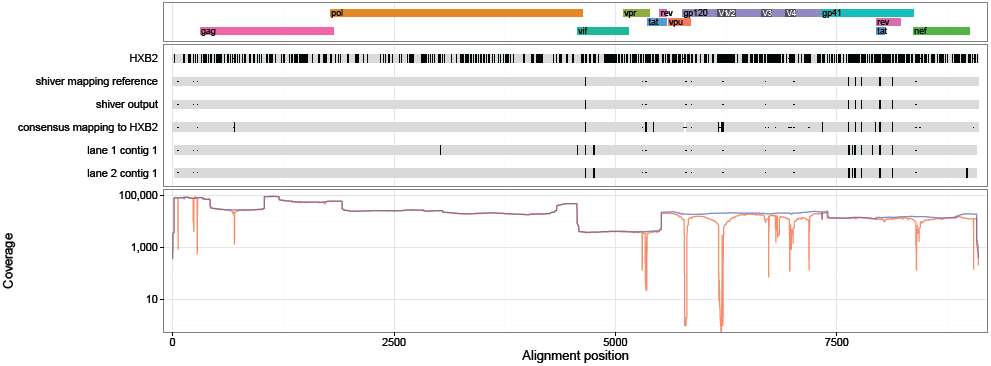
19960_3_49 sequences and coverage (mapping to the `shiver` reference in blue, to HXB2 in red).

**Figure 115:**
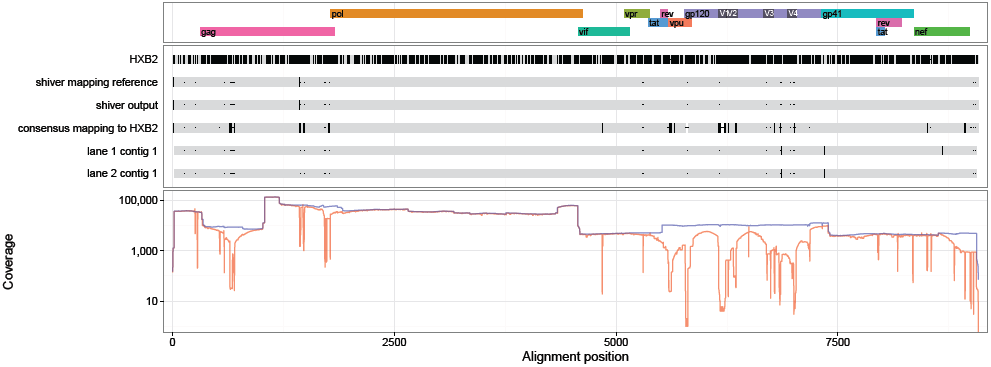
19960_3_6 sequences and coverage (mapping to the `shiver` reference in blue, to HXB2 in red).

**Figure 116:**
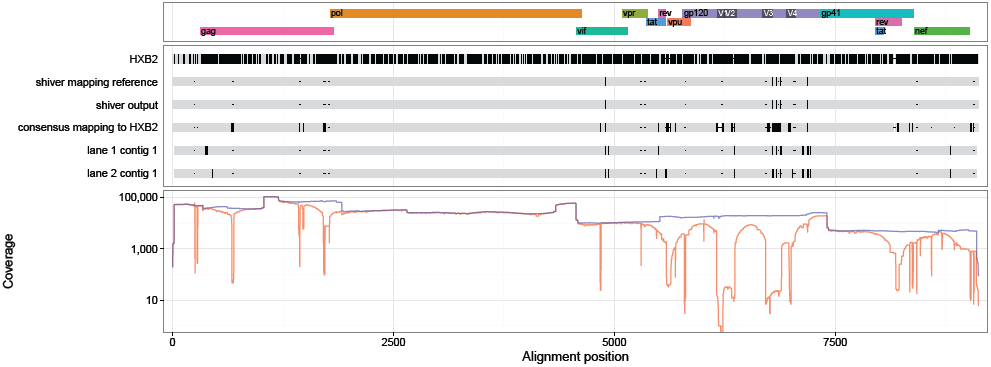
19960_3_70 sequences and coverage (mapping to the `shiver` reference in blue, to HXB2 in red).

**Figure 117:**
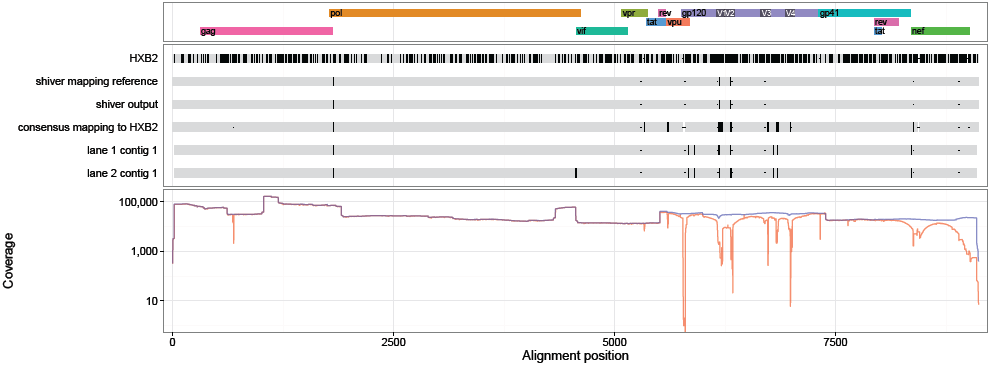
19960_3_9 sequences and coverage (mapping to the `shiver` reference in blue, to HXB2 in red).

**Figure 118:**
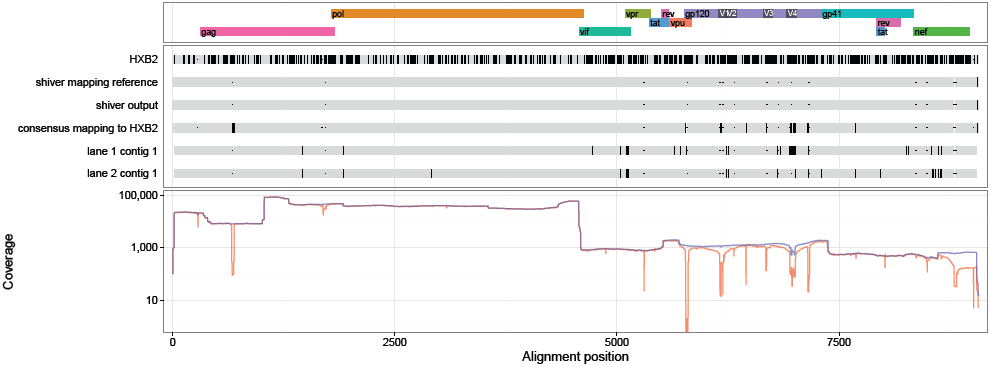
20004_3_146 sequences and coverage (mapping to the `shiver` reference in blue, to HXB2 in red).

**Figure 119:**
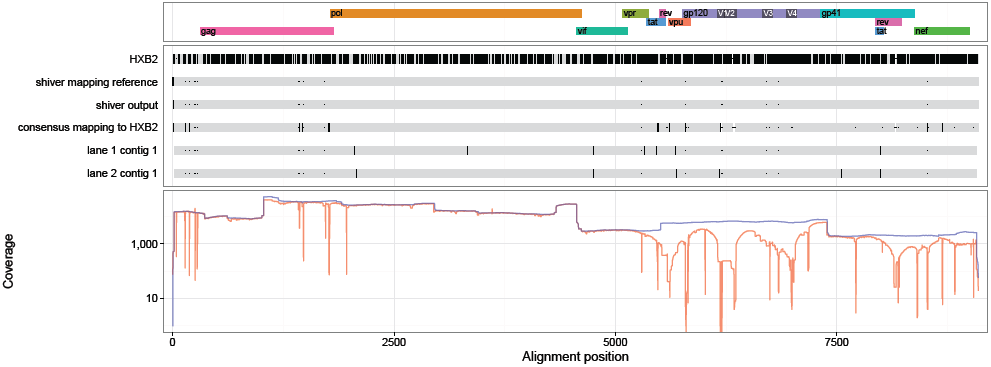
20004_3_155 sequences and coverage (mapping to the `shiver` reference in blue, to HXB2 in red).

**Figure 120:**
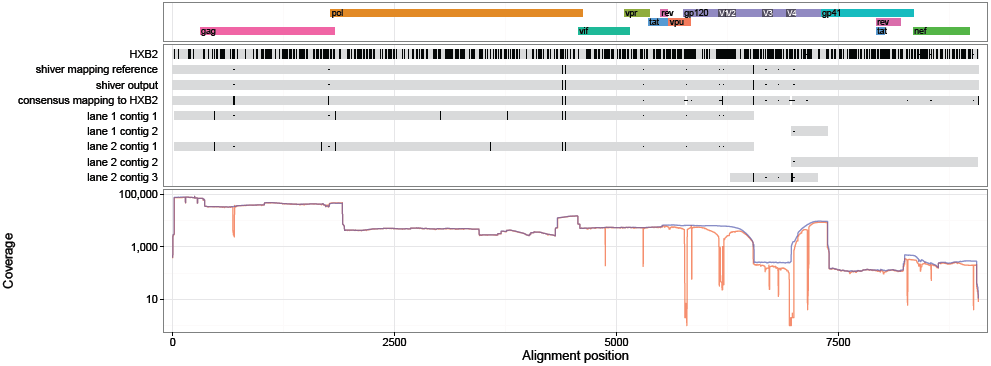
20004_3_56 sequences and coverage (mapping to the `shiver` reference in blue, to HXB2 in red).

